# Odorant receptors for floral- and plant-derived volatiles in the yellow fever mosquito, *Aedes aegypti* (Diptera: Culicidae)

**DOI:** 10.1101/2023.10.17.562234

**Authors:** Heidi Pullmann-Lindsley, Robert Huff, John Boyi, R. Jason Pitts

## Abstract

Adult mosquitoes require regular sugar meals, usually floral nectar, to survive and flourish in natural habitats. Both males and females locate potential sugar sources using sensory proteins called odorant receptors activated by plant volatiles that facilitate orientation toward flowers or honeydew. The Yellow Fever mosquito, *Aedes aegypti* (Linnaeus, 1762), possesses a large repertoire of odorant receptors, many of which are likely to support floral odor detection and nectar-seeking. In this study, we have employed a heterologous expression system and the two-electrode voltage clamping technique to identify environmentally relevant chemical compounds that activate specific odorant receptors. Importantly, we have uncovered ligand-receptor pairings for a suite of *Aedes aegypti* odorant receptors likely to mediate appetitive or aversive behavioral responses, thus shaping a critical aspect of the life history of a medically important mosquito. Moreover, the high degree of conservation of these receptors in other disease-transmitting species suggests common mechanisms of floral odor detection. This knowledge can be used to further investigate mosquito foraging behavior to either enhance existing, or develop novel, control strategies, especially those that incorporate mosquito bait-and-kill or attractive toxic sugar bait technologies.

## Introduction

Arthropods rely on sensory receptors to interpret and interact with their environments. Arguably, olfaction is the most important sensory modality, conveying the ability to sense volatile chemical cues over large distances [1–3]. Olfaction is responsible for many arthropod behaviors, including host-seeking [4, 5], oviposition [6, 7], mate selection [8, 9], and nectar foraging [10, 11]. Odorant receptors (ORs) comprise the most prominent family of volatile odor detectors encoded in mosquito genomes [12]. Another family of proteins called ionotropic receptors (IRs) also functions in insect olfaction, especially in organic acid detection [13, 14]. ORs are transmembrane proteins expressed by odorant receptor neurons housed in hairlike projections called sensilla [15, 16]. Sensilla sensory appendages such as antennae, maxillary palps, labella, and leg tarsi constitute the primary olfactory appendages in insects [17, 18, 19]. ORs are ligand-gated ion channels that respond to diverse, environmentally relevant volatile odorants [20]. ORs most likely form multimers that include co-expression of a single type of tuning OR complexed with an odorant receptor coreceptor (Orco), which is highly conserved across most insect taxa [21–23]. Chemical volatiles enter the sensilla through tiny pores on their surfaces and either diffuse through the sensilla lymph or are shuttled by odorant binding proteins to the neuronal membrane, where they can interact with the OR/Orco complex [24, 25]. Generally, single ORs respond to one compound most robustly, though they may demonstrate the responsiveness of a handful of other ligands. Most arthropod species have between 50 and 100 different ORs, which allow for complex and robust interactions with their respective environments through the perception of various volatile compounds. However, some species have as few as five or as many as 300 [26, 27]. The diversity of receptor proteins underlies distinct behavioral phenomena across arthropod species.

OR characterization is a crucial step toward arthropod control and surveillance. By delving into the molecular intricacies of how these receptors function, we can significantly contribute to our understanding of insect behavior. Deorphanizing single ORs has yielded valuable insights into many insect species, directly informing behavioral control strategies. One of the most comprehensively studied relationships lies in the realm of pheromones [28]. In the silk moth (*Bombyx mori),* an OR, BmOR1, that responded to the female-produced pheromone bombykol was identified [29, 30] and characterized using a heterologous expression system [31]. Notably, disrupting BmOR1 led to the complete loss of sensitivity to bombykol, subsequently impacting pheromone source detection in male moths[32]. While the intricate relationships between receptors, ligands, and behavior are best characterized in the context of pheromone reception, ORs are linked to numerous other behaviors, including nectar foraging. Understanding their function could prove invaluable to combatting arthropod-borne threats.

The prominent disease vector *Aedes aegypti* (Linnaeus, 1762) has 133 putative odorant receptors that vary across sex, life stage, and tissue [33–35]. *Ae. aegypti* is a prominent disease vector can transmit several arboviruses, including Dengue, Zika, and yellow fever. This species inhabits much of Africa, Asia, South America, Oceania, and parts of North America and Europe [36]. However, as climate change continues to alter ecosystems and global urbanization accelerates the viable range of *Ae. aegypti* may also expand, and its threat to human health could increase [37, 38]. Despite the significant impact of this species, relatively few ORs have been characterized, specifically ones presumed to contribute to female host-seeking behaviors [39, 40]. Single sensilla recordings (SSR) have revealed that *Ae. aegypti* neurons responded to a variety of chemical compounds collected from human volunteers, including octanal and nonanal [41]. We have also previously demonstrated that an OR highly conserved in Culicidae disease vectors responds to carboxylic acids present on human skin [5]. However, other essential behaviors likely mediated by ORs have yet to be studied. Remarkably, little is known about the shared behavior of nectar foraging between males and females, which is crucial for mosquito survival and reproductive fitness [42]. While preference between nectar sources has been documented in numerous mosquito species, the molecular and olfactory underpinnings are yet to be elucidated [43–45]. This behavior is of particular importance for disease control as characterizing the ORs that mediate it may provide vital information about potential nectar-seeking linked attractants that can be used in mosquito bait and kill (MBAK) or attractive toxic sugar bait (ATSB) systems [46–48]. While MBAKs and ATSBs have demonstrated relative success in reducing mosquito populations [49, 50], pursuing a more specific attraction to mosquito species could reduce off-target effects [51].

In this study, we investigated ORs in *Ae. aegypti* that are conserved in several disease vector mosquito species and are closely related to other ORs that function in floral odor sensing. These odorant receptors are expressed mainly in male and female antennae [33–35], suggesting they may function in a shared behavior such as nectar foraging. We initiated this study by completing manual gene annotations and intron analysis [52]. Following this, we used a heterologous expression system and a technique called two-electrode voltage clamping (TEVC) to identify essential ligands [53]. After synthesizing cRNA for odorant receptors of interest, we injected unfertilized *Xenopus laevis* oocytes with the cRNA for the OR of interest (called the tuning receptor) and cRNA for *Ae. aegypti* Orco. After allowing time for protein expression, we used TEVC to record changes in membrane potential as a library of 147 potential ligands was perfused past. Using this method, we identified probable ligands for ten ORs, eight of which were novel or revealed a previously unknown ligand. Many odorant receptors responded to indole-like compounds that affect *Ae. aegypti* behavior [54, 55]. These studies highlight the importance of characterizing ORs to improve our understanding of the molecular basis for *Ae. aegypti* nectar seeking or other behaviors.

While we have demonstrated that these receptors and ligands interact on a molecular basis, more research is needed to determine how the ligands might influence a mosquito’s behavior. Future studies will include olfactometry with wild-type and gene-knockout mosquitoes to verify the role of these ORs and research into the expression patterns of these genes in response to ligand exposure. These studies can help to identify specific attractants or repellants for use in surveillance systems or bait traps to help control mosquito populations and combat the spread of disease.

## Materials and Methods

### Phylogenetic Analysis and Gene Annotations

A conserved clade of genes appears to be activated by floral or nectar odors and utilized by mosquitoes in foraging. These genes were identified via BLAST via tBLASTn or BLASTp using VectorBase and confirmed using available RNAseq data (S1 Figure). Closely related ORs from *Aedes albopictus* (Skuse, 1894) and *Culex quinquefasciatus* (Say, 1823) were also identified. DNA sequences from the reference genomes (*Ae. aegypti* LVP_AGWG, *Ae. albopictus* Foshan FPA, *Cx. quinquefasciatus* JHB 2020) were downloaded and the genes were manually annotated using SnapGene (Dotmatics, Boston, MA, USA). Intron analysis was conducted for the *Ae. aegypti* genes and compared with previously identified conserved introns [52]. A Neighbor-Joining tree based on amino acid alignments was constructed using Geneious Prime software (Biomatters Ltd., Boston, MA, USA).

### Gene cloning and Sequencing

The coding regions of ten genes of interest (S2 Table) and AaegOrco were synthesized by Twist Bioscience (San Francisco, CA, USA) and cloned into the pENTR plasmid. The Gateway LR directional cloning system (Invitrogen Corp., Carlsbad, CA, USA) was utilized to subclone AaegOR coding regions into the pSP64t-RFA oocyte expression plasmid. Plasmids were purified using the GeneJet Plasmid Miniprep Kit (ThermoFisher Scientific, Waltham, MA, USA). Purified plasmids were sequenced in both directions to confirm coding regions.

### OR Gene Expression Analysis and de novo Transcript Assembly

OR expression values in Transcripts Per Million (TPM) were obtained from VectorBase (vectorbase.org) based on published studies of *Ae. aegypti* chemosensory appendages [34]. *De novo* transcripts were assembled using non-bloodfed female antennae and maxillary palp RNAseq data sets [34] using the Trinity software package [35] on the Galaxy server (usegalaxy.org, S3 Figure).

### Chemical Reagents

Chemical compounds (S4 Table) used in the deorphanization of AaegORs were purchased from Acros Organics (Morris, NJ, USA), Alfa Aesar (Ward Hill, MA, USA), and ThermoFisher Scientific (Waltham, MA, USA) at the highest purity available. Compounds were stored as 1M stocks in 100% DMSO. Serial dilutions of each compound were prepared in 10% DMSO and blended by chemical class.

### Two-electrode voltage clamping

pSP64t-RFA: AaegOR clones were linearized using Xbal, and cRNAs were synthesized using the mMESSAGE mMACHINE ® SP6 kit (Life Technologies, Carlsbad, CA, USA). Xenopus laevis stage V-VII oocytes purchased from Xenopus1 (Dexter, MI, USA) were stored at 18°C in Incubation medium (ND96 96 with 5% dialyzed horse serum,50μg/mL gentamycin,100μg/mL streptomycin, 100μg/mL penicillin, and 550μg/mL sodium pyruvate). 30nL of cRNA was injected into multiple oocytes using a microinjection syringe pump (World Precision Instruments, Sarasota, FL, USA) maintained at 18C for 48-72 hrs. Odorant blends were perfused across individual oocytes for 10-15 seconds, held at a resting membrane potential of -80 mV using an OC-725C oocyte clamp (Warner Instruments, LLC, Hamden, CT, USA). Induced currents were measured indirectly using the two-electrode voltage clamp technique (TEVC). Following each stimulus, currents were allowed to return to baseline before further stimulation. Data was captured using the Digidata 1550 B digitizer and PClamp10 software (Molecular Devices, San Jose, CA, USA). Concentration-response curves were generated by perfusing unitary odorants over oocytes at increasing concentrations from [10^-3^M] to [10^-9^M]. Data were analyzed using GraphPad Prism 8 (GraphPad Software Inc., La Jolla, CA, USA). At least two biological replicates were conducted at each concentration and for each compound. Data for at least seven oocytes responding to each compound was analyzed (S6 File).

## Results

### 1. *Aedes aegypti* odorant receptors are expressed in male and female antennae and conserved in other disease vector mosquitoes

Analysis of publicly available RNAseq data sets [36] demonstrated that the ORs in our study are expressed in adult sensory appendages (Table 1). With the exception of AaegOR8, which is expressed in the maxillary palps [33], ORs were most abundant in the antennae of both sexes (Table 1; [34]). Phylogenetic analysis of possible floral volatile sensing odorant receptors in three major disease vectors revealed several different groupings, which may indicate shared or similar ligands (Figure 1A). *Aedes aegypti* ionotropic receptor 75d (AaegIR75d) served as an outgroup. Available RNAseq analysis [36] confirmed gene structures (S1 Figure). Our analysis revealed that there are homologs to the *Ae. aegypti* genes of interest in both *Ae. albopictus* and *Cx. quinquefasciatus* and manual annotations revealed close homology (S5 Figure) and highly conserved intron structures (Figure 1B). Phylogenetic analysis supports homologies between ORs from multiple vector species, providing targets for future functional studies, as the receptors deorphanized in this study are likely to. Through our analysis, we identified and named six new ORs in *Ae. albopictus.* We located two OR11 homologs, named AalbOR11a and AalbOR11b, as well as a secondary OR13 homolog (AalbOR13b), and three previously unannotated OR31 homologs (AalbOR31a, AalbOr31b, AalbOR31c). As with the previously annotated ORs, these clustered in the cladogram with the homologous genes from *Ae. aegypti* and *Cx. quinquefasciatus*.

**Figure 1.**
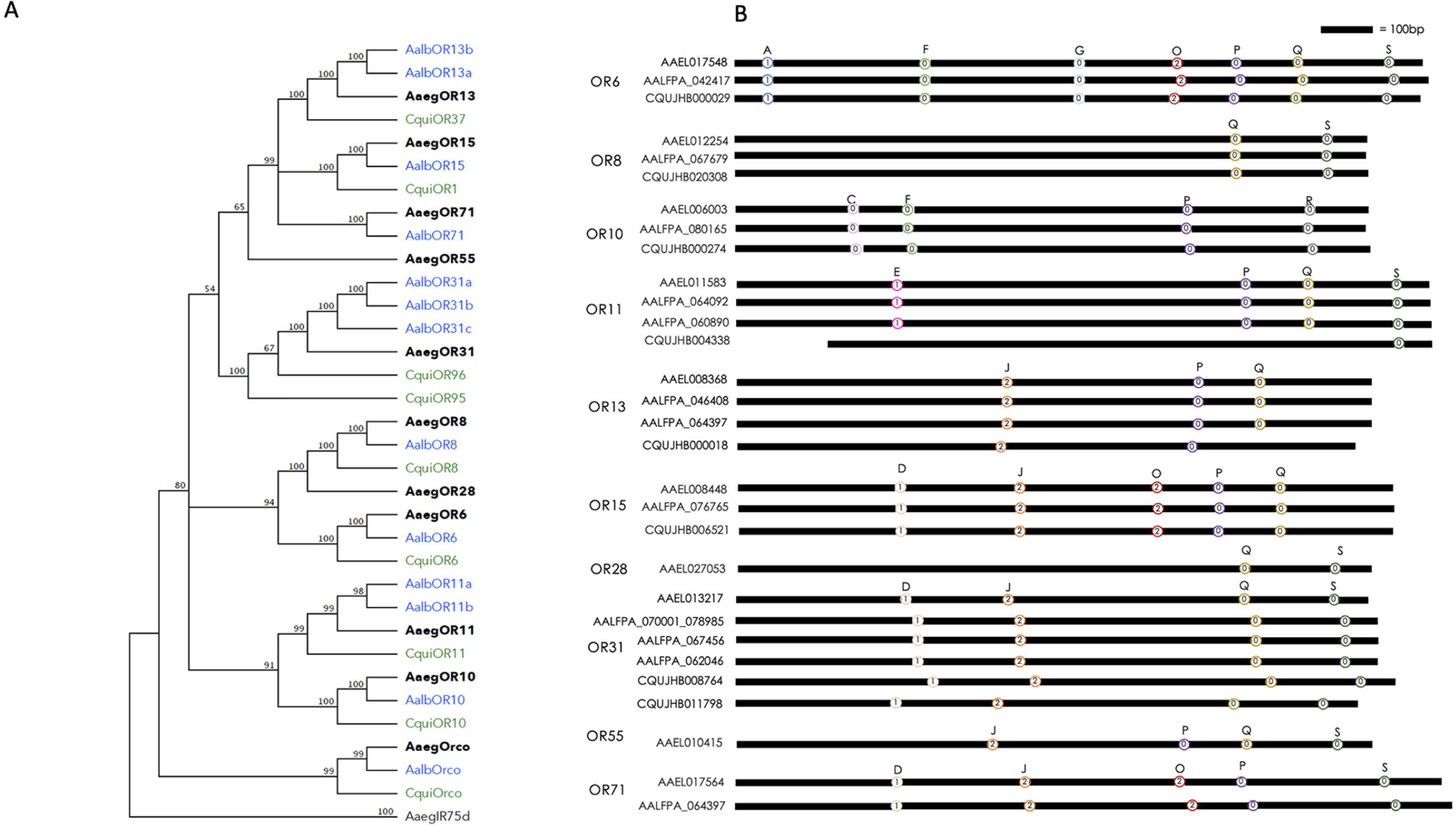
Phylogenetic analysis of possible floral volatile sensing ORs in major disease vectors (A) Neighbor-joining tree showing relationships of floral volatile sensing OR conceptual peptides for three major disease vector species of mosquitoes *Ae. aegypti* (black), *Ae. albopictus* (Blue), and *Cx. quinquefasciatus* (green). Node values signify bootstrap values from 1000 pseudoreplicates. (B) Intron analysis indicates shared exon and intron sizes and structures. Letter notation indicates intron, and colors vary based on likely designation.

**Table 1:**
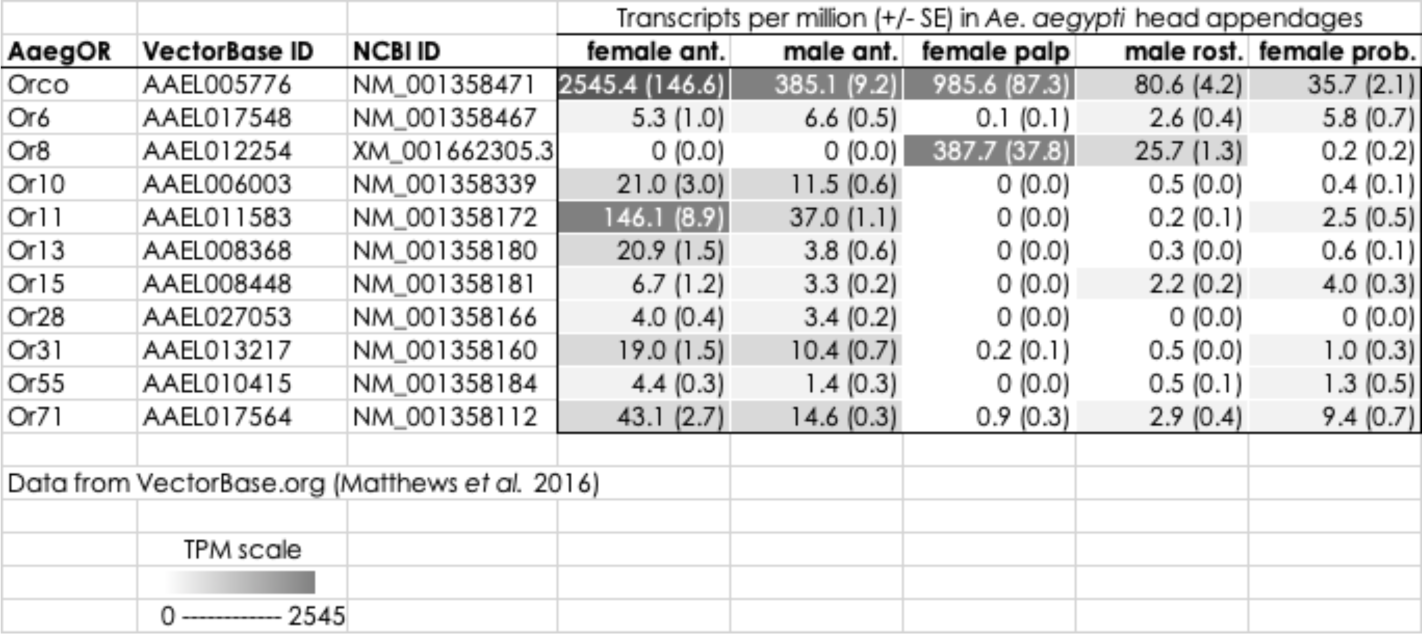
Abundance of AaegOR transcripts in *Ae. aegypti* sensory appendages.

### 2. Odorant receptors respond to environmentally relevant compounds in a concentration dependent manner

*Ae. aegypti* ORs responded to blends of chemical compounds (Figure 2). While there is some responsiveness to multiple blends by specific ORs, all demonstrated a pronounced specificity for one blend over the others, as indicated by highly positive kurtosis values (Table 2). The blends that most frequently activated ORs were indole or alcohol blends. Unitary compounds from the highest activating blend for each OR were tested individually. We determined the primary ligand each OR responded to from our chemical library and noted other secondary ligands (Table 2, Figure 3). As with the blends, ORs exhibited pronounced responses to a single ligand, again supported by highly positive kurtosis values. A defining feature of OR responsiveness is that it is concentration-based; we also investigated responses to increasing concentrations with that compound. We determined the half-maximal effective concentration (EC_50_) for each primary ligand (Table 2, Figure 4).

**Figure 2.**
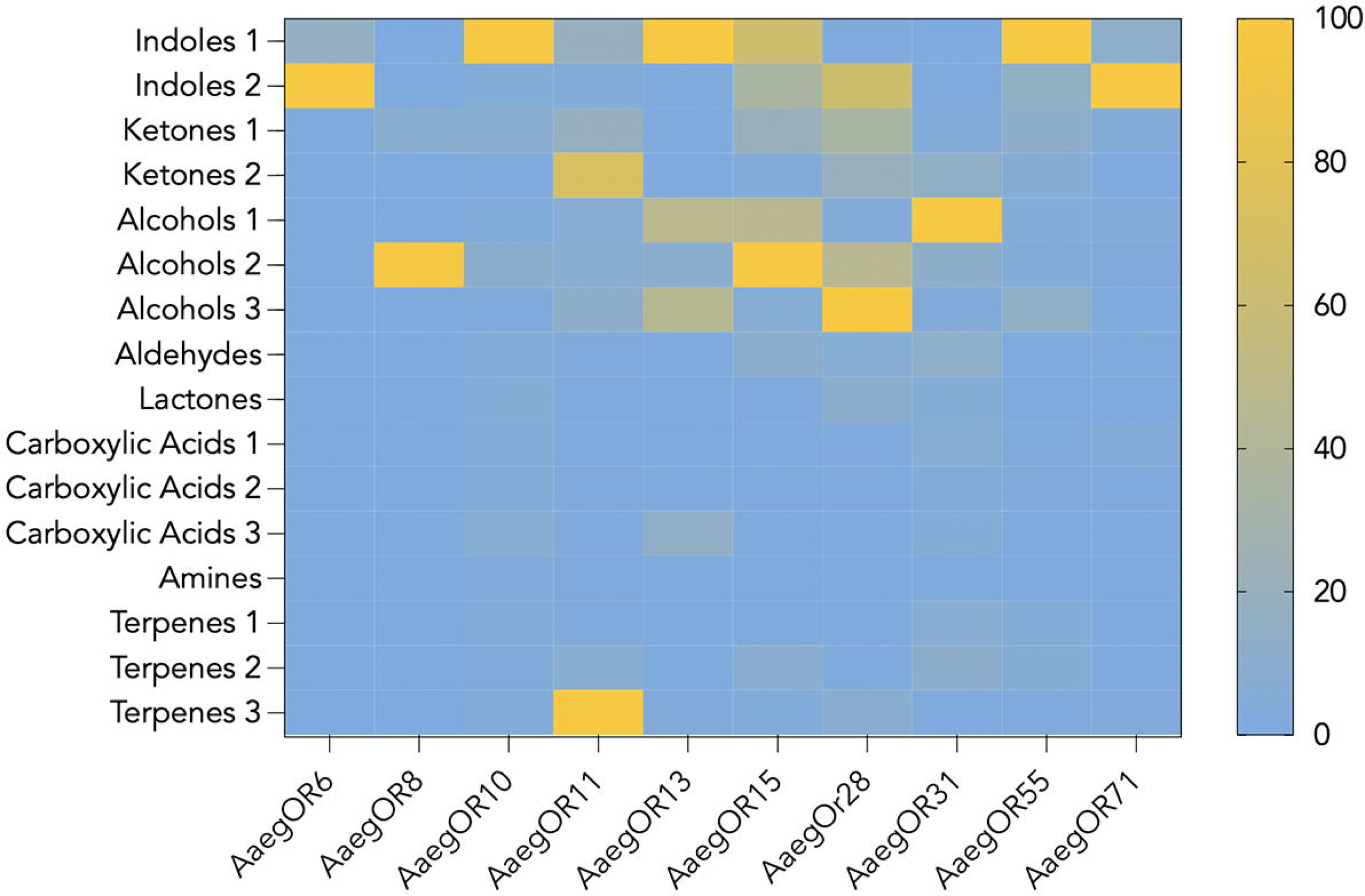
Heat map depicting AaegOR responses to blends of environmentally relevant odors. Blends of chemical compounds used for receptor screening are listed on the lefthand side of the figure while receptor identities are listed below. Color gradation is shown on the right and represents normalized mean oocyte responses from highest (100, yellow) to lowest (0, blue).

**Figure 3.**
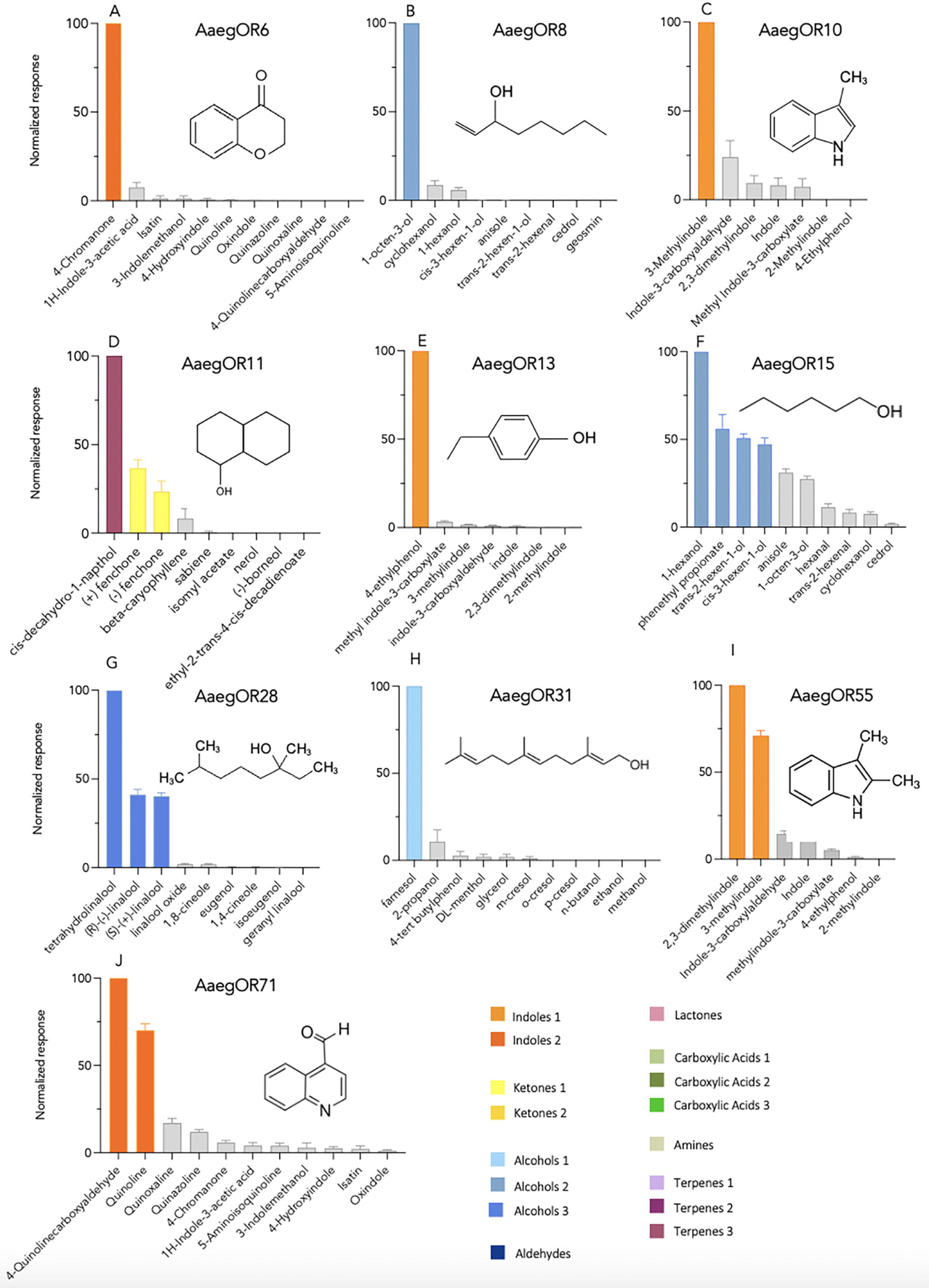
*Ae. aegypti* odorant receptor responses to unitary compounds. Normalized mean current responses of AaegORX + AaegOrco expressing oocytes (n=7–10 oocytes per blend) to the individual components of various blends (A) 4-chromanone (B) 1-octen-3-ol, (C) 3-methylindole (D) cis-decahydro-1-napthol, (E) 4-ethyl-phenol, (F) 1-hexanol, (G) tetrahydrolinalool, (H) farnesol, (I) 2,3-dimethylindole, and (J) 4-quinolinecarboxyaldehyde. The structure of each compound is shown in figure inserts.

**Figure 4.**
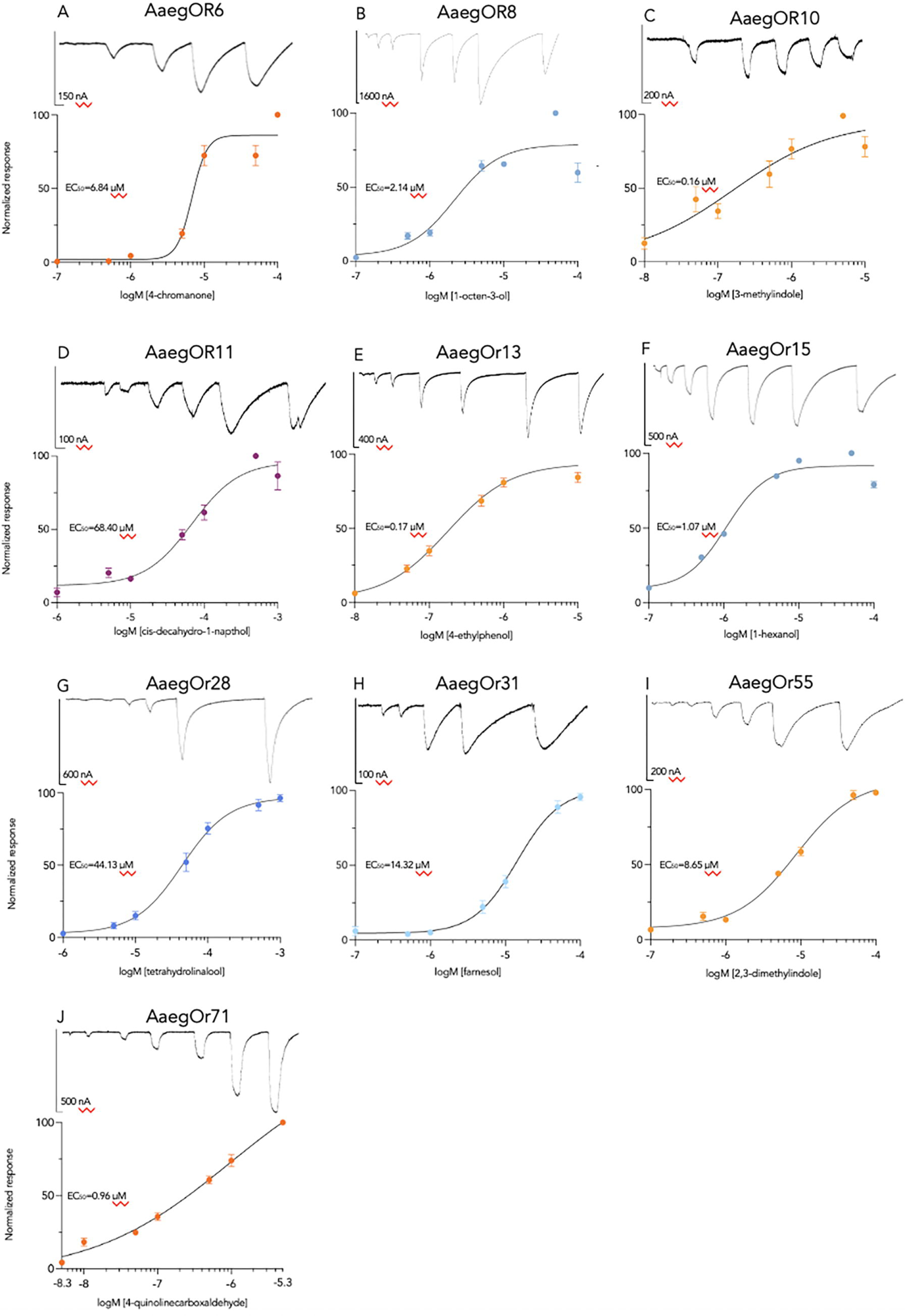
*Ae. aegypti* odorant receptor concentration response curves. Normalized mean current responses of AaegORX + AaegOrco expressing oocytes (n>10 per concentration) to (A) 4-chromanone, (B) 1-octen-3-ol, (C) 3-methylindole (D) cis-decahydro-1-napthol, (E) 4-ethyl-phenol, (F) 1-hexanol, (G) tetrahydrolinalool, (H) farnesol, (I) 2,3-dimethylindole, and (J) 4-quinolinecarboxyaldehyde. Representative traces from oocyte recordings are shown above each curve. EC50 values are shown in figure insets.

**Table 2:** Functional responses of *Ae. aegypti* odorant receptors.

| Odorant Receptor | Blend | Kurtosis | Primary Ligand (s) | Kurtosis | EC<sub>50</sub> (μM) | Secondary Ligand(s) |
| --- | --- | --- | --- | --- | --- | --- |
| AaegOr6 | Indoles 2 | 14.86 | 4-chromanone | 10.85 | 6.84 |  |
| AaegOr8 | Alcohols 2 | 15.71 | 1-octen-3-ol | 8.75 | 2.14 |  |
| AaegOr10 | Indoles 1 | 15.61 | 3-methylindole | 5.87 | 0.16 |  |
| AaegOr11 | Terpenes 3 | 5.68 | cis-deca-hydro-1-naphthol | 6.85 | 68.40 | fenchone |
| AaegOr13 | Indoles 1 | 6.24 | 4-ethyl-phenol | 9.11 | 0.17 |  |
| AaegOr15 | Alcohols 2 | 3.63 | 1-hexanol | 5.23 | 1.07 | phenethyl propionate |
| AaegOr28 | Alcohols 2 | 15.96 | tetrahydrolinalool | 7.70 | 44.13 |  |
| AaegOr31 | Alcohols 1 | 14.58 | Farnesol | 10.70 | 14.32 |  |
| AaegOr55 | Indoles 1 | 14.50 | 2,3-dimethylindole | 0.22 | 8.65 |  |
| AaegOr71 | Indoles 2 | 15.56 | 4-quinolinecarboxaldehyde | 3.08 | 0.96 |  |

Summarization table of 10 *Ae. aegypti* odorant receptors and the corresponding responsiveness to chemical and unitary compound blends. Kurtosis values are indicated for both blend and unitary compounds. EC50 values (µM) are indicated for primary ligands. Secondary ligands are presented if identified during the functional characterizations.

Three ORs responded most robustly to the Indole 1 blend. OR10, OR13, and OR55 responded to 3-methylindole (0.16 µM), 4-ethylphenol (0.17 µM), and 2,3-dimethylindole (8.65 µM), respectively. OR6 responded to 4-chromanone (6.81 µM) from the Indole 2 blend. The other receptor that responded to this blend was OR71, which responded to 4-quinoline carboxaldehyde (0.96 µM). From the Alcohols 1 blend, OR31 responded to farnesol (14.32 µM), while from the Alcohols 2 blend, OR8 and OR15 responded to 1-octen-3-ol (2.14 µM) and 1-hexanol (1.07 µM), respectively. A previously identified ligand of OR15, phenethyl propionate, demonstrated an EC_50_ of 20.16 µM. OR28 responded to tetrahydrolinalool (44.13 µM) from the Alcohols 3 blend. Finally, the only receptor to respond to a compound not belonging to an indole or alcohol blend was OR11, which responded to cis-decahydro-1-naphthal (68.40 µM) from the Terpenes 3 blend. We found that OR11 has an EC_50_ of about 77.10 µM to a previously identified ligand, fenchone.

## Discussion

For mosquitoes, nectar foraging to obtain sugar meals is essential for survival. Individuals that can locate and access higher-quality nectar sources exhibit extended lifespans and increased fecundity [43, 57–58]. Therefore, exploring the molecular mechanisms driving selective preferences is imperative. While past research has demonstrated the existence of nectar source preferences in various mosquito species corresponding to nectar qualities [44–45, 59–63], a substantial gap in our comprehension of the sensory components facilitating these preferences still needs to be addressed. In other insect species, the study of ORs has proven instrumental in deciphering behavioral patterns and formulating effective control measures [64–66].

ORs function as sensory gateways, providing critical information about the surrounding environment and enabling mosquitoes to discern and select nectar sources based on the chemical signals they receive [3]. In this context, our research centers on the responsiveness of a series of ORs to a library of ecologically relevant compounds, linking OR-ligand relationships to floral odor sensing. By expanding our knowledge of the chemoreceptive basis of a critical mosquito behavior, we expect to contribute to enhanced surveillance and control measures by developing more effective odor-baited trap and kill systems. In turn, these improvements may help to mitigate the impact of mosquitoes on human health.

We began this study by examining *Ae. aegypti* OR gene structures, expression patterns, and relatedness to ORs from two other prominent disease vectors, *Cx. quinquefasciatus* and *Ae. albopictus* [67]. We selected a subset of ORs to characterize based on their homologies to ORs or that have been previously shown to respond to floral odors [68]. While AaegORs are expressed in various chemosensory tissues, antennae are the primary structures involved in odor sensing [1]. Their expression in the antennae of both sexes suggests that ORs are used for shared behaviors, such as nectar feeding. Furthermore, by refining and examining *Ae. aegypti* ORs gene structures, we identified homologs of interest in both *Ae. albopictus* and *Cx. quinquefasciatus,* two medically relevant mosquito species. The existence of numerous closely related genes leads us to propose that the essential act of nectar foraging in vector mosquitoes shares a common underlying OR mechanism, indeed that OR-ligand pairs may have been conserved through divergence of these species. Future studies will be required to characterize the response profiles of conserved ORs, and to establish the requirement of these receptors in nectar foraging in *Ae. aegypti* and other species

Using two-electrode voltage clamping in conjunction with the *Xenopus laevis* oocyte expression system, we assessed the responses of the AaegORs to our chemical library, which contains 147 environmentally significant compounds organized into 14 blends. Many AaegORs responded robustly to the blends containing indole or indole-like compounds (Supp. Table 1). Indoles are relatively ubiquitous in natural settings and are present in potential nectar sources. AaegOR71 responded to 4-quinoline carboxaldehyde, a compound identified in honey samples and endophytic bacteria via gas chromatography [69–70]. However, indoles are emitted naturally from many sources, including flowers, vertebrate hosts, and oviposition sites of microbiotic origin, suggesting they influence multiple behaviors across species [71–72]. AaegOR6 responds best to 4-chromanone, a compound that is found in shrubs of the Asteraceae family, is structurally related to quinolines and is a key constituent of flavonoid skeletons [73–74]. AaegOR10 responds to 3-methylindole (skatole) found in animal feces and produces behavioral responses in other Dipteran species [75–78]. 4-ethylphenol, the primary ligand of AaegOR13, is a volatile constituent in wine and animal waste and has been linked to oviposition in *Culex pipiens* [78–79]. While there is little evidence about natural sources of 2,3-dimethylindole, the AaegOR55 ligand is closely related structurally to 3-methylindole. Other ORs responded to compounds from the “Alcohols 1”, “Alcohols 2”, or “Alcohols 3” blends. AaegOR8 responds to 1-octen-3-ol, a well-characterized fungal volatile and insect attractant [80]. A green-leaf volatile, 1-hexanol, is another compound with well-characterized physiological effects in many arthropods [81–84]. Our study discovered that 1-hexanol activates AaegOR15 with a low threshold (EC_50_ ∼1.07 µM). AaegOR28 responds to tetrahydrolinalool, which is structurally related to linalool, another plant-derived ligand to mosquito ORs [85]. AaegOR31 responds to farnesol, a significant constituent of the honeybee Nasanov pheromone [86]. Lastly, the AaegOR11 ligand, cis-decahydro-1-napthol, is structurally related to geosmin, a known oviposition attractant [87]. One limitation of oocyte studies is the number of compounds that can be screened. More efficacious or potent ligands may exist for AaegOR11 and AaegOR28, which produced EC_50_ values greater than 40µM (Table 2). We will continue to expand our library by adding compounds structurally related to known ligands.

While four (AaegOR8, AaegOR10, AaegOR11, and AaegOR15) of the ORs tested in this study already have identified ligands, we assessed them to uncover more potentially behaviorally relevant ligands, as our chemical library differs from others tested. For AaegOR8 and AaegOR10, our data confirm previous observations of high selectivity to 1-octen-3-ol and skatole, respectively [56, 88]. For AaegOR11 and AaegOR15, our data supports and adds to previous studies. For AaegOR11, we found that cis-decahydro-1-napthol produced a greater magnitude of response than (-)-fenchone or (+)-fenchone, and we identified 1-hexanol as a potent activator of AaegOR15 in addition to phenethyl propionate. This demonstrates that even for deorphanized ORs, assessing their function with expanded libraries can aid in further understanding receptor-ligand interactions.

*Ae. aegypti* and other vector mosquitoes remain persistent global threats to human health and flourishing. Pursuing receptor-ligand pairings is paramount, as it opens doors to identifying potential attractants for behavioral control or targets for genetic knockout. In our investigation of floral-odor sensing receptors, we aim to pinpoint compounds that impart behavioral attraction for both sexes. Such compounds can be applied in Mosquito Baited Traps (MBAK) and Attractive Toxic Sugar Baits (ATSB) systems. While these technologies have been successfully deployed in some studies [48–52], identifying compounds that activate *Ae. aegypti* ORs offers the potential to refine and enhance their effectiveness while minimizing the impact on non-target species. OR deorphanization is a crucial initial step in unraveling the intricate sensory systems of arthropods, providing valuable insights for future experiments and control strategies. This focused endeavor not only aids in addressing the immediate threats posed by vector mosquitoes but also contributes to our broader understanding of insect behavior and the development of sustainable control measures.

## Supporting Information

S1 Figure. **RNA sequencing validation of ORs**

Pictures showing RNA abundance and gene structure of selected ten (10) Odorant receptors (Or) present in unique antennal and maxillary palp reads from Matthews *et al.,* 2018 gotten from Vectorbase. The scale on the left represents the values of transcripts per million. Highlighted in red and blue are the exons present in each Or. The actual abundance of RNA per exon is represented in orange. Numbers at the top of the figures represent chromosome numbers and positions within the AaegL5 assembly. Vectorbase gene ID and descriptions are present in black and blue text on the pictures.

S2 Table. **Names and peptide sequences for *Ae. aegypti* ORs and homologs**

Table lists names, VectorBase identification, *Ae. aegypti* homolog, and name attribution

S3 Figure. **Sequence alignments of *de novo* transcripts and ORs**

Transcripts were generated using RNA seq data from Matthews *et al.,* 2018 [35] and *Aedes aegypti* transcripts (AaegL5) of the ten (10) Odorant receptors (OR). Sequences have been aligned using ClustalW with character counts (Li *et al.,* 2015). Gene IDs of the AaegL5 transcripts and those of the denovo transcripts are shown on the left-hand side. Identical residues are highlighted in gray while different residues are highlighted in black. Missing residues are not highlighted

S4 Table. **List of chemical compounds used in TEVC**

S5 Figure. **Homolog alignments and intron locations**

Alignments between *Ae. Aegypti, Ae. Albopictus,* and *Cx. quinquefasciatus* odorant receptors were completed using Clustal Omenga (https://www.ebi.ac.uk/Tools/msa/clustalo/). Gray shading indicated shared residues, green shading indicated intron position.

## Acknowledgements

This work was supported by a grant from the National Institute of Allergy and Infectious Diseases (award # 1R01AI148300-01A1 to RJP). The authors also acknowledge Drs. Jeffrey Riffell (University of Washington) and Omar Akbari (University of California San Diego) and members of their respective laboratories for helpful discussions of results. The authors are grateful to the Baylor University Molecular Biosciences Center (Dr. Michelle Nemec, Director) for the use of various equipment and resources.

