## Supplementary figures and images for "Odorant receptors for floral- and plant-derived volatiles in the yellow fever mosquito, *Aedes aegypti* (Diptera: Culicidae)"

### Supplemental File 1

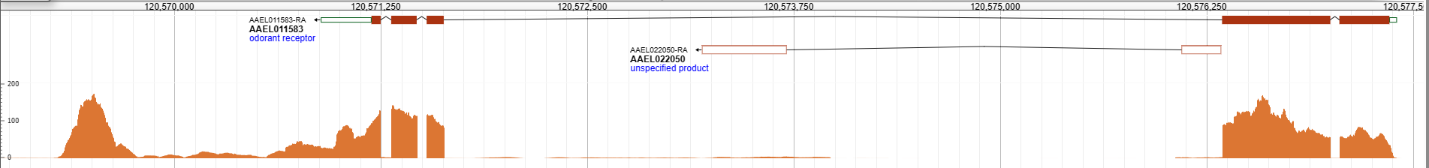

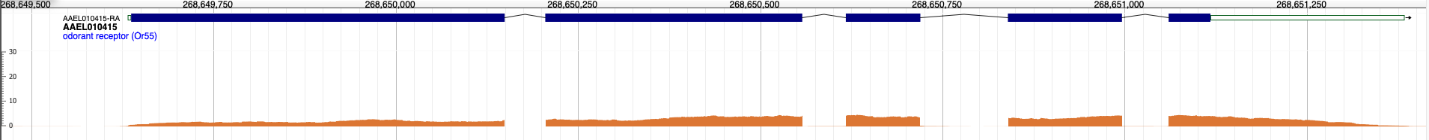

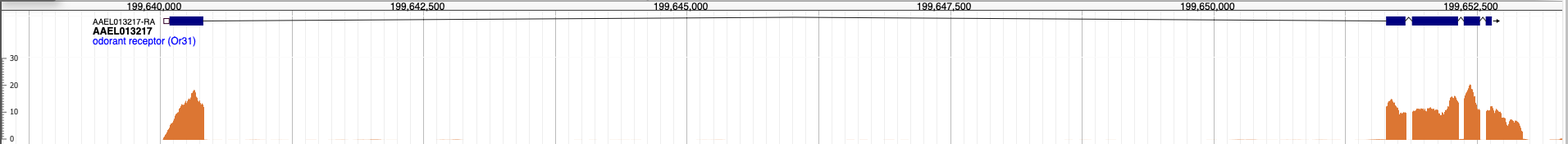

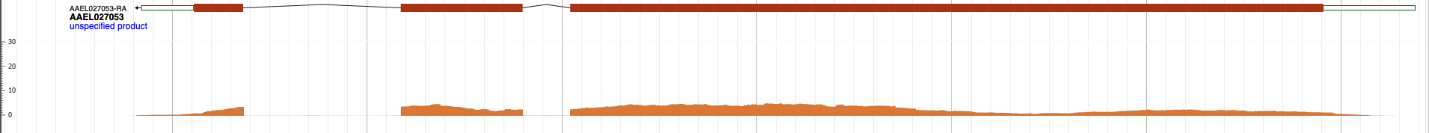

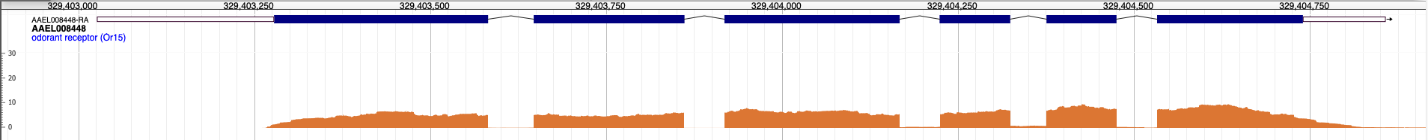

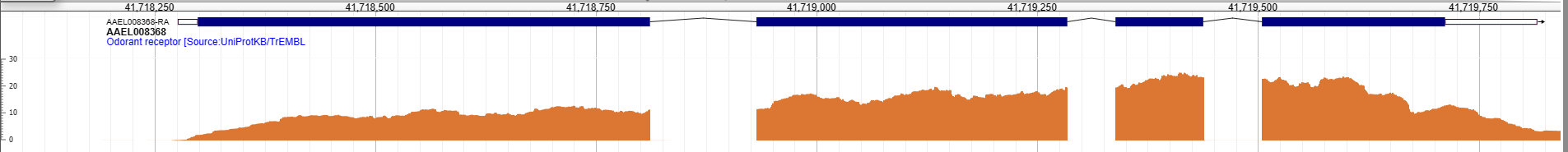

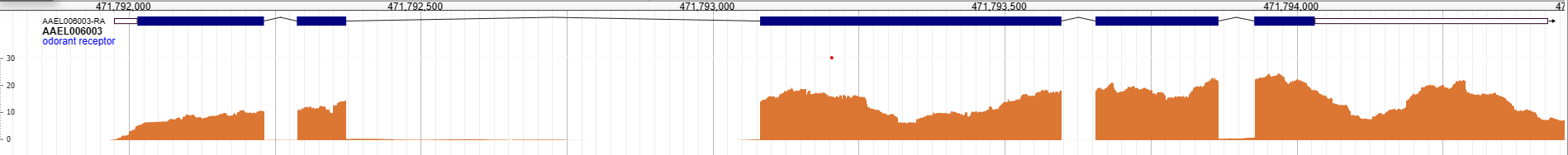

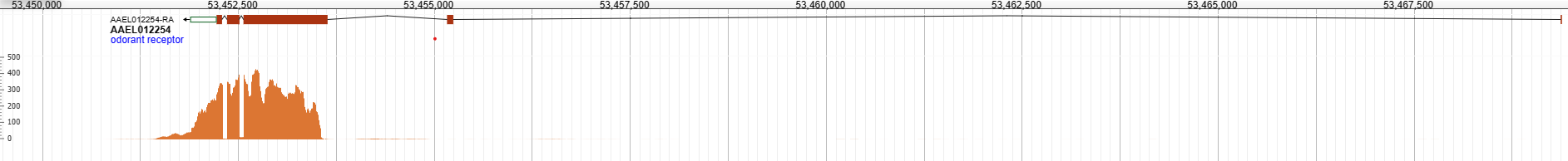

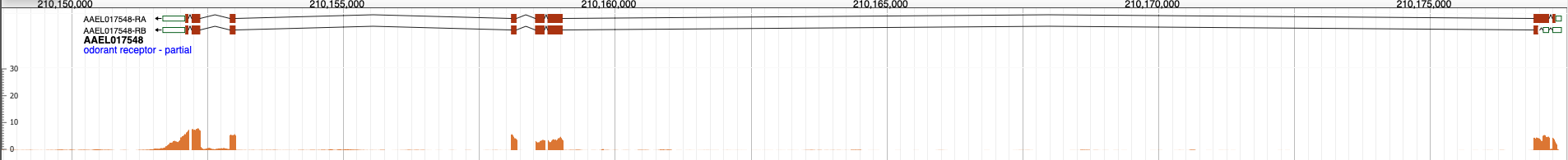

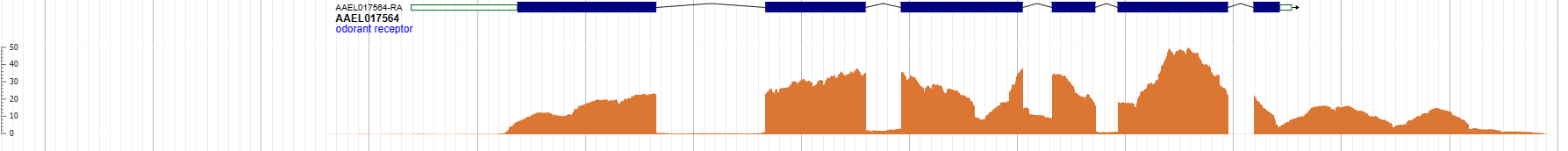
