## Supplemental File 3 for "Odorant receptors for floral- and plant-derived volatiles in the yellow fever mosquito, *Aedes aegypti* (Diptera: Culicidae)"

```
AaegOr10_AAEL006003-RA                           MASILDCPIVSVNARVWRFWSFVLKHDAMRYISIIPVTVMTFFMFTDLCRSWGNIQELII	60
AaegOr10_Antenna_TRINITY_DN1780_c0_g1_i10        MASILDCPIVSVNARVWRFWSFVLKHDAMRYISIIPVTVMTFFMFTDLCRSWGNIQELII	60


AaegOr10_AAEL006003-RA                           KAYFAVLYFNAVLRTLILVKDRKLYENFMQGISNVYFEISHIDDHKIQSLLKSYTVRARM	120
AaegOr10_Antenna_TRINITY_DN1780_c0_g1_i10        KAYFAVLYFNAVLRTLILVKDRKLYENFMQGISNVYFEISHIDDHKIQSLLKSYTVRARM	120


AaegOr10_AAEL006003-RA                           LSISNLALGAIISTCFVVYPIFTGERGLPYGMFIPGLDSFRSPHYEIIYIVQVVLTFPGC	180
AaegOr10_Antenna_TRINITY_DN1780_c0_g1_i10        LSISNLALGAIISTCFVVYPIFTGERGLPYGMFIPGLDSFRSPHYEIIYIVQVVLTFPGC	180


AaegOr10_AAEL006003-RA                           CMYIPFTSFFASTTLFGLVQIKTLQRQLQTFKDNISSQDKEKVKTKVVKLIEDHKRIITY	240
AaegOr10_Antenna_TRINITY_DN1780_c0_g1_i10        CMYIPFTSFFASTTLFGLVQIKTLQRQLQTFKDNINSQDKEKVKAKVVKLIEDHKRIITY	240


AaegOr10_AAEL006003-RA                           VSELNSLVTYICFVEFLSFGMMLCALLFLLNVIENHAQIVIVAAYIFMIISQIFAFYWHA	300
AaegOr10_Antenna_TRINITY_DN1780_c0_g1_i10        VSELNSLVTYICFVEFLSFGMMLCALLFLLNVIENHAQIVIVAAYIFMIISQIFAFYWHA	300


AaegOr10_AAEL006003-RA                           NEVREESMNLAEAAYSGPWVELDNSIKKKLLLIILRAQQPLEITVGNVYPMTLEMFQSLL	360
AaegOr10_Antenna_TRINITY_DN1780_c0_g1_i10        NEVREESMNLAEAAYSGPWVELDNSIKKKLLLIILRAQQPLEITVGNVYPMTLEMFQSLL	360


AaegOr10_AAEL006003-RA                           NASYSYFTLLRRVYN	375
AaegOr10_Antenna_TRINITY_DN1780_c0_g1_i10        NASYSYFTLLRRVYN	375


AaegOr11_AAEL011583-RA                          MQLKDEWIQEDDVYDNPLLRLTINGLKYYGILLYKSQPFKKLNCFRGVCFTASMLAFNVT	60
AaegOr11_Antenna_TRINITY_DN1275_c0_g1_i2        MQLKDEWIQEDDVYDNPLLRLTINGLKYYGILLYKSQPFKKLNCFRGVCFTASMLAFNVT	60


AaegOr11_AAEL011583-RA                          QYVDLYQVWGNIAEMTANAATTLLFTTTIVRILHFYWNRARFNNAIKVADEGVQHLLRFG	120
AaegOr11_Antenna_TRINITY_DN1275_c0_g1_i2        QYVDLYQVWGNIAEMTANAATTLLFTTTIVRILHFYWNRARFNNAIKVADEGVQHLLRFG	120


AaegOr11_AAEL011583-RA                          NAPEKEIFWDNVKYMNRLTAAFWICALVTANTMCVYALVQYQSLKSMDSFNSTEPFDPPT	180
AaegOr11_Antenna_TRINITY_DN1275_c0_g1_i2        NAPEKEIFWDNVKYMNRLTAAFWICALVTANTMCVYALVQYQTLKSIDSFNSTEPFDPPT	180


AaegOr11_AAEL011583-RA                          ILRSWYPTDNIVDSFATIYLIQLYIMYVGQLIVPCWHVFMVSLMLYARTALMALNYKLAN	240
AaegOr11_Antenna_TRINITY_DN1275_c0_g1_i2        ILRSWYPTDNIVDSFATIYLIQLYIMYVGQLIVPCWHVFMVSLMLYARTALMALNYKLAH	240


AaegOr11_AAEL011583-RA                          LEQYAVSGMRSGRKIKCVVDPEEERCNVRKELIVECIQQQHKIFEYTRELEALTRGAMFM	300
AaegOr11_Antenna_TRINITY_DN1275_c0_g1_i2        LEQYAVSGIRSGRKIKCVVDPEEERCNVRKELIVECIQQQHKIFEYTRELEALTRGAMFM	300


AaegOr11_AAEL011583-RA                          DFVVFSVLLCALLFEASSTNSFVQIFIDICYIMTMTAMLFLYYWHANEIHYQANLLSSSA	360
AaegOr11_Antenna_TRINITY_DN1275_c0_g1_i2        DFVVFSVLLCALLFEASSTNSFVQIFIDICYIMTMTAILFLYYWHANEIHYQANLLSSSA	360

AaegOr11_AAEL011583-RA                          FMNDWYNYPRSVNRHLITFICYSNKPLDMKAYIVSMSLDTFLAILRASYSYFTILKQAAG	420
AaegOr11_Antenna_TRINITY_DN1275_c0_g1_i2        FMNDWYNYPRSVNRHLITFICYSNKPLDMKAYIVSMSLDTFLAILRASYSYFTILKQAAG	420


AaegOr13_AAEL008368-RA                          MWQPLRKFLAPGPELLSFGLQMLRFIGLWGDRRQVVRYLLVLFSEFIFLIGPKALLGSDK	60
AaegOr13_Antenna_TRINITY_DN9791_c2_g1_i2        MWQPLRKFLAPGPELLSFGLQMLRFIGLWGDRRQVVRYLLVLFSEFIFLIGPKALLGSDK	60


AaegOr13_AAEL008368-RA                          EGFDSTARNIGELIFLVEVCISIGIFASRRASFERLIVVLENILRRKWPRNLQDEIYRFH	120
AaegOr13_Antenna_TRINITY_DN9791_c2_g1_i2        EGFDSTARNIGELIFLVEVCISIGIFASRRASFERLIVVLENILRRKWPRNLQDEIYRFH	120


AaegOr13_AAEL008368-RA                          RRMEFFARAYALYIGFLLFLYNCVPIGSTIVKLIRFDESERSDFMLVVELQFFWFDIRRN	180
AaegOr13_Antenna_TRINITY_DN9791_c2_g1_i2        RRMEFFARAYALYIGFLLFLYNCVPIGSTIVKLIRFDESERSDFMLVVELQFFWFDIRRN	180


AaegOr13_AAEL008368-RA                          VVHYAIYMAFCFVAVSCSAYQSTLKGSVIVVVTQYGSKLFELISKRIDAMRSIERAADRD	240
AaegOr13_Antenna_TRINITY_DN9791_c2_g1_i2        VVHYAIYMAFCFVAVSCSAYQSTLKGSVIVVVTQYGSKLFELISKRIDAMRSIERAADRD	240


AaegOr13_AAEL008368-RA                          RELREIVKLHGMALEYVQHLESTISFVMINQIMNCIFIWCLMMFYVSTNFGPNAANVMLL	300
AaegOr13_Antenna_TRINITY_DN9791_c2_g1_i2        RELREIVKLHGMALEYVQHLESTISFVMINQIMNCIFIWCLMMFYVSTNFGPNAANVMLL	300


AaegOr13_AAEL008368-RA                          FLVLMGEMVVYCLNGTALSEQAAGVGHAIYNYPWYKESVAMQKNMQLMIQRAQRPTGITA	360
AaegOr13_Antenna_TRINITY_DN9791_c2_g1_i2        FLVLMGEMVVYCLNGTALSEQAAGVGHAIYNYPWYKESVAMQKNMQLMIQRAQRPTGITA	360


AaegOr13_AAEL008368-RA                          AKFYFVNIERLGLVTQASYSYYLILKNRF	389
AaegOr13_Antenna_TRINITY_DN9791_c2_g1_i2        AKFYFVNIERLGLVTQASYSYYLILKNRF	389


AaegOr15_AAEL008448-RA                         MKYFELTEPEAAMPLALRLLETYGLRGGKRKFLQFQVTILWELLMIVIPKIVFGYRSQDL	60
AaegOr15_Antenna_TRINITY_DN448_c0_g1_i4        --HFELTEPEAAMPLALRLLETYGLRGGKRKFLQFQVTILWELLMIVIPKIVFGYRSQDL	58


AaegOr15_AAEL008448-RA                         VIRGLSELLFQLHIMIRISIFAWHRFKYESLIDIIRKVYRKTFSTGGDPTSKSIILKFNQ	120
AaegOr15_Antenna_TRINITY_DN448_c0_g1_i4        VIRGLSELLFQLHIMIRISIFAWHRFKYESLIDIIRKVYRKTFSTGGDPTSKSIILKFNQ	118


AaegOr15_AAEL008448-RA                         MINKQSKGYFLYIMGCVSLFSVAPVVQSVIIFMANQSRNGTEKAEYVTMMEQEFYGLDIR	180
AaegOr15_Antenna_TRINITY_DN448_c0_g1_i4        MINKQSKGYFLYIMGCVSLFSVAPVVQSVIIFMANQSRNGTEKAEYVTMMEQEFYGLDIR	178


AaegOr15_AAEL008448-RA                         GNFGHYAIYVALAGLAHYYSASFFAVTGVIIICGVRCTILTFKLINVRLSKLHELPKQDI	240
AaegOr15_Antenna_TRINITY_DN448_c0_g1_i4        GNFGHYAIYVALAGLAHYYSASFFAVTGVIIICGVRCTILTFKLINVRLSKLHELPKQDI	238


AaegOr15_AAEL008448-RA                         RDELREIIDLHVDALRCIQLLEQIANLAMVIQIIDCVLIWISMILYMRNNLGVDAISLMV	300
AaegOr15_Antenna_TRINITY_DN448_c0_g1_i4        RDELREIIDLHVDALRCIQLLEQIANLAMVIQIIDCVLIWISMILYIRNNLGVDAISLMV	298


AaegOr15_AAEL008448-RA                         LFVALTGETYALCDLLTQLTSESLAVTRAIIDCQWYSLPLDVQKSLSFVLFRAQRKEGIT	360
AaegOr15_Antenna_TRINITY_DN448_c0_g1_i4        LFVALTGETYALCDLLTQLTSESLAVTRAIIDCQWYSLPLDVQKSLSFVLFRAQRKEGIT	358


AaegOr15_AAEL008448-RA                         AAKFFFMDIERFGSVAQTSYSIYVVLKDQL	390
AaegOr15_Antenna_TRINITY_DN448_c0_g1_i4        AAKFFFMDIERFGSVAQTSYSIYVVLKDQL	388


AaegOr28_AAEL027053-RA                           MVWIEADRSSSLEYDSFFRLPKIFGLLNGVVYNDEKPSSKWSKAKNVYFWISLMHSILVA	60
AaegOr28_Antenna_TRINITY_DN41586_c0_g1_i1        MVWIEADRSSSLEYDSFFRLPKIFGLLNGVVYNDEKPSSKWSKAKNVYFWISLMHSILVA	60


AaegOr28_AAEL027053-RA                           VLELVYLAKSVEQNADFVFIMSLVPLVGHGILAIVKLSVQKYYHKEINSILISLKGIYPS	120
AaegOrO28_Antenna_TRINITY_DN41586_c0_g1_i1       VLELVYLAKSVEQNADFVFIMSLVPLVGHGILAIVKLSVQKYYHKEINSILISLKGIYPS	120


AaegOr28_AAEL027053-RA                           TLDDTITKDYSKKILYMKLFVIFYLVTLIFFNIVPFAPVLHTYFTTGVFEKTLPFFIYYW	180
AaegOr28_Antenna_TRINITY_DN41586_c0_g1_i1        TLDDTITKDYSKKILYMKLFVIFYLVTLIFFNIVPFAPVLHTYFTTGVFEKTLPFFIYYW	180


AaegOr28_AAEL027053-RA                           YDWRRPILYELTFIEQIWVSTASVVANMNIDLMLCSLILQISMHFDVLSDRLSVLQHNDH	240
AaegOr28_Antenna_TRINITY_DN41586_c0_g1_i1        YDWRRPILYELTFIEQIWVSTASVVANMNIDLMLCSLILQISMHFDVLSDRLSVLQHNDH	240


AaegOr28_AAEL027053-RA                           KELAKCVERHSVLLDLCLRVENIFSRSMLASFLLSSVIICLTGFQVFAQDSINKAIPYAT	300
AaegOr28_Antenna_TRINITY_DN41586_c0_g1_i1        KELAKCVERHSVLLDLCLRVENIFSRSMLASFLLSSVIICLTGFQVFAQDSINKAIPYAT	300


AaegOr28_AAEL027053-RA                           FLFLHMVDVYLLCYYGNLMMEKSLAVSNYAYESLWYLGNRPFQKSILIILERGQRAQKLT	360
AaegOr28_Antenna_TRINITY_DN41586_c0_g1_i1        FLFLHMVDVYLLCYYGNLMMEKSLDVSNYAYESLWYLGNRPFQKSILIILERGQRAQELT	360


AaegOr28_AAEL027053-RA                           AMKFIDINLTCFKTILSTSFSYFTLLKVLNEPME	394
AaegOr28_Antenna_TRINITY_DN41586_c0_g1_i1        AMKFIVINLTCFKTILSTSFSYFTLLKVLNEPME	394


AaegOr31_AAEL013217-RA                          MAPTQNGRDREKFLRVQLLCLALIGIKRHETVSSRTIFHVCFISMVIMDLATILFALEHA	60
AaegOr31_Antenna_TRINITY_DN6991_c0_g1_i1        MAPTQNGRDREKFLRVQLLCLALIGIKRHETVSSRTIFHVCFISMVIMDLATILFALEHA	60


AaegOr31_AAEL013217-RA                          NDIALVCDCLGPTFTAYLGIVKQYCLSAHRVELWNIIETLRRLKDYAGISEIESIERNNK	120
AaegOr31_Antenna_TRINITY_DN6991_c0_g1_i1        NDIALVCDCLGPTFTAYLGIVKQYCLSAHRVELWNIIETLRRLKDYAGTSEIESIERNNK	120


AaegOr31_AAEL013217-RA                          IDRFLATAYLMSASATGSLFIIAALAKGCYKLIFQNIIEWGFPLSLSFPFKTSHPIVFGM	180
AaegOr31_Antenna_TRINITY_DN6991_c0_g1_i1        IDRFLATAYLMSASATGSLFIIAALAKGCYKLIFQNIIEWGFPLSLSFPFKTSHPIVFGV	180


AaegOr31_AAEL013217-RA                          FFVWSSAAIYIVVFCSVSSDASFGGLASNVVVHFKLLQKRLQDATFADNDENLKQLIEYH	240
AaegOr31_Antenna_TRINITY_DN6991_c0_g1_i1        FFVWSSAAIYIVVFCSVSSDASFGGLASNVVVHFKLLQKRLQDATFADNDENLKQLIEYH	240


AaegOr31_AAEL013217-RA                          SLLLNLSRKIMSSFRVIIINNLLVASVLLCVLGFQLVMFLGSTLMLIYLMYVTAIVIQIT	300
AaegOr31_Antenna_TRINITY_DN6991_c0_g1_i1        SLLLNLSRKIMSSFRVIIINNLLVASVLLCVLGFQLVMFLGSTLMLIYLMYVTAIVIQIT	300


AaegOr31_AAEL013217-RA                          FFAYYGSLLLHESEEVSISIYCSNWYEASPKTRRILLQCLMRAQVPVNTKAGFMVASLPT	360
AaegOr31_Antenna_TRINITY_DN6991_c0_g1_i1        FFAYYGSLLSHESEEVSSSIYCSNWYEASPKTRRILLQCLMRAQVPVNTKAGFMVASLPT	360


AaegOr31_AAEL013217-RA                          LRAILNSAGSYVALLLSFTDN	381
AaegOr31_Antenna_TRINITY_DN6991_c0_g1_i1        LRAILNSAGSYVALLLSFTDN	381


AaegOr55_AAEL010415-RA                           MQQKPPFRLMSASLKLCRWLGLWHDVNLDKPCWQTVFISFCLLFWYILPGCLYITRGERM	60
AaegOr55_Antenna_TRINITY_DN13286_c0_g1_i6        -NSQPPFRLMSASLKLCKWLGLWHDANLDKPCWQTVFLILCLLFWYILPGYMYIVRGEKM	59


AaegOr55_AAEL010415-RA                           LQYLLKSILEVFSMCVIVLRCVVHMINRKTVQNCFVELQDAISTFENSPYEDVRQMLRHL	120
AaegOr55_Antenna_TRINITY_DN13286_c0_g1_i6        LQDLLKPILEVFSMAVIVLRCLIHMINRQSVQECFADLQNAISKFKNSPYEDVQKILRHL	119


AaegOr55_AAEL010415-RA                           LKSADYLVKIYVSIVFIQASIYGLVPAILTTYRYCNSNETIQLPSAVMEADYVLFDHSTN	180
AaegOr55_Antenna_TRINITY_DN13286_c0_g1_i6        LKSADYIVKFYVSIVFVQAGLYGFLSAALTTFKYCTSNEIIQLPSAVMDADYVLFDHTVN	179


AaegOr55_AAEL010415-RA                           YWIWLLVTIISLLVEYLLLGTFSSQECLFWNLLHHVSSLLKVVRLEIARLDQYTDPKQYT	240
AaegOr55_Antenna_TRINITY_DN13286_c0_g1_i6        YWIWLPVTIVSLIIEYLMLGSISAQECLFWNLLHHISSLFKIVHLEIARLDQYKDPKQFK	239


AaegOr55_AAEL010415-RA                           ERLASIVSTHEVCYRCARSLEIVLSPLLAVLYCTCITQTCYLLFVISMIDDLVVIASMIF	300
AaegOr55_Antenna_TRINITY_DN13286_c0_g1_i6        ERLAFIVSIHEVCFRSARCMEKVLSPLLALWYCACIAQTCYLLFFISMVNDVVVVASMIF	299


AaegOr55_AAEL010415-RA                           VLQYIVFLIFSFSMLGAELTEESALVSEAIYNSNWYMRMPAERRLLLFMKMRADRPVGIT	360
AaegOr55_Antenna_TRINITY_DN13286_c0_g1_i6        VLQYVVFLIFSFSMLGAELMEESARVSDAVYNSNWYMRMPAERRLLLFMKMRADRPVGIT	359


AaegOr55_AAEL010415-RA                           AAKFFYVNRSTFAEAMKTAFSFFTIMQQFYGE	392
AaegOr55_Antenna_TRINITY_DN13286_c0_g1_i6        AAKFFYVNRSTFAEAMKTAFSFFTIMQ-----	386


AaegOr71_AAEL017564-RA                            MELSYHRSLPPELQIMPFQLRCMELIGLIGPKGRFYRFVLAFGWGTFVILLPKSVLGIGS	60
AaegOr71_Antenna_TRINITY_DN13495_c0_g1_i23        MELSYHRSLPPELQIMPFQLRCMELIGLIGPKGRFYRFVLAFGWGTFVILLPKSVLGIGS	60


AaegOr71_AAEL017564-RA                            SELDAIIKGFAELLFEGNLFIAVASLVPKLPLVKRLLHVLSEIFRQATHDKHDAKDHCYA	120
AaegOr71_Antenna_TRINITY_DN13495_c0_g1_i23        SELDAIIKGFAELLFEGNLFIAVASLVPKLPLVKRLLHVLSEIFRQATHDKRDVKDHCYA	120


AaegOr71_AAEL017564-RA                            LICEQNSKIDKFCKFYFIYCCFGPFVFCIPAMVTSYVRYFDTTNNGTNANGSEHLRFELP	180
AaegOr71_Antenna_TRINITY_DN13495_c0_g1_i23        LICEQNSKIDKFCKFYFIYCCFGPFVFCIPAMVTSYVRYFGTTNNGTNANGSEHLRFELP	180


AaegOr71_AAEL017564-RA                            MEQEFYWLPIRTNFACYHLFLALSLSAYCVCSYMSVIKVSTLLIMIKYCSLVYRLVAIRI	240
AaegOr71_Antenna_TRINITY_DN13495_c0_g1_i23        MEQEFYWLPIRTNFACYHLFLALSLSAYCVCSYMSVIKVSTLLIMIKYCSLVYRLVAIRI	240


AaegOr71_AAEL017564-RA                            RELGKXPPGRKQDDTDEDERTKMVMVKEVVEMHEKALEATDLVEKVINIPIAMQFMACIL	300
AaegOr71_Antenna_TRINITY_DN13495_c0_g1_i23        RELGK*PPGRKQDDTDEDERTKMVMVKEVVEMHEKALEATDLVEKVINIPIAMQFMACIL	299


AaegOr71_AAEL017564-RA                            FWCMTMVYVSTNINFNLFNVMVLFWLSLIETYGYSYLGTELSEDAKAVGHAVYDLPWYED	360
AaegOr71_Antenna_TRINITY_DN13495_c0_g1_i23        FWCMTMVYVSTNINFNLFNVMVLFWLSLIETYGYSYLGTELSEDAKAVGHAVYDLPWYED	359


AaegOr71_AAEL017564-RA                            SAQLQRYYRLMIQRSQQNIGVTAAKFFIVGIEKFGKVVNLSYSYYLVLKDVLDRL	415
AaegOr71_Antenna_TRINITY_DN13495_c0_g1_i23        SAQLQRYYRLMIQRSQQNIGVTAAKFFIVGIEKFGKVVNLSYSYYLVLKDVLDRL	414


AaegOr6_AAEL017548-RA                           MVSFEQSFTAVDLILITSGIPSCSSFYPVSIKSALKRNAVFLMAFTLLFYTAFGELVYLV	60
AaegOr6_Antenna_TRINITY_DN29927_c0_g2_i1        ------------------------------------------------------------	0


AaegOr6_AAEL017548-RA                           EMLQRDFSFLEITFQAPCLGYCTIGLMKMLILAVKRNTIAELVQSLQEEWNKSVRSLEHQ	120
AaegOr6_Antenna_TRINITY_DN29927_c0_g2_i1        ------------------------------------------------------------	0


AaegOr6_AAEL017548-RA                           SICDDVMKPAIRFTTIVAVVNIVMGLAFTLLPIPEMIYYYSTHGKWVRQLPFLIWWSFDA	180
AaegOr6_Antenna_TRINITY_DN29927_c0_g2_i1        ------------------------------------------------------------	0


AaegOr6_AAEL017548-RA                           YSGFVYYFIYPLYVVIGFSGIIIHMGFDCLFCILSAHLCVHLRILKFDLENLTYGLDSSE	240
AaegOr6_Antenna_TRINITY_DN29927_c0_g2_i1        ------------------------------------------------LENLTYGLDSSE	12


AaegOr6_AAEL017548-RA                           NMSIKLNPKLFYIVEKHQNILECHDRMNKIFNFALFYNFFVSSFIICIQGFMVTAASGYT	300
AaegOr6_Antenna_TRINITY_DN29927_c0_g2_i1        NMSIKLNPKLFYIVEKHQNILECHDRMNKIFNFALFYNFFVSSFIICIQGFMVTAASGYT	72


AaegOr6_AAEL017548-RA                           LIKFALFLASFLVELFLLCFYGHHIVESSVLVAEAAYNCLWYNTNHQFRTIILQMVNKGQ	360
AaegOr6_Antenna_TRINITY_DN29927_c0_g2_i1        LIKFALFLASFLVELFLLCFYGHHIVESSVLVAEAAYNCLWYNTNHQFRTIILQMVNKGQ	132


AaegOr6_AAEL017548-RA                           IPLSLMAWKIWPVNMNTFANILSASWSYFTLIRTVYAD	398
AaegOr6_Antenna_TRINITY_DN29927_c0_g2_i1        IPLSLMAWKIWPVNMNTFANILSASWSYFTLIRTVYAD	170


AaegOr8                                    MNDLVKFESFIRVPEIFFDMIGITRYGEARDTWKARLKQAFFWSSYANTIFCLIIEHIYF	60
AaegOr8_palp_TRINITY_DN10600_c0_g1_i2      MNDLVKFESFIRVPEIFFDMIGITRYGEARDTWKARLKQAFFWSSYANTIFCLIIEHIYF	60


AaegOr8                                    IKAAGNFTNFLELTALAPCIGFTALSIVKIMTIKLNEAKLNGILDRLSDLFPRSHLDQDR	120
AaegOr8_palp_TRINITY_DN10600_c0_g1_i2      IKAAGNFTNFLELTALAPCIGFTALSIVKIMTIKLNEAKLNGILDRLSDLFPRSHLDQDR	120


AaegOr8                                    YRTYNYNLESQMVMKSFSILYMILIWIFNLLPLVSMLVNYISTGILEKELPYFMWYWYDW	180
AaegOr8_palp_TRINITY_DN10600_c0_g1_i2      YRTYNYNLESQMVMKSFSILYMILIWIFNLLPLVSMLVNYISTGILEKELPYFMWYWYDW	180


AaegOr8                                    HKAGYYEITFFHQNWGAFDSAVFNLSTDLLFCAIILLICLQFDILAYRLRHAKGDYKELE	240
AaegOr8_palp_TRINITY_DN10600_c0_g1_i2      HKAGYYEITFFHQNWGAFDSAVFNLSTDLLFCAIILLICLQFDILAYRLRHAKGDYKELE	240


AaegOr8                                    QCVKLHQSVVELSNQLEGIFSPSILVNFVGSSVIICLVGFQATSNISAFDLFKFILFLIS	300
AaegOr8_palp_TRINITY_DN10600_c0_g1_i2      QCVKLHQSVVELSNQLEGIFSPSILVNFVGSSVIICLVGFQATSNISAFDLFKFILFLIS	300


AaegOr8                                    SLVQVFLLCYYGNKLIEASSQIGYCAFEGTWYMADLRYQKSLLFVMTRAGQWQKLTAMKF	360
AaegOr8_palp_TRINITY_DN10600_c0_g1_i2      SLVQVFLLCYYGNKLIEASSQIGYCAFEGTWYMADLRYQKSLLFVMTRAGQWQKLTAMKF	360


AaegOr8                                    SVVSLASYSAILSTSFSYFTLLKTIYEPSQK	391
AaegOr8_palp_TRINITY_DN10600_c0_g1_i2      SVVSLASYSAILSTSFSYFTLLKTIYEPSQK	391
```
