## Supplemental File 5 for "Odorant receptors for floral- and plant-derived volatiles in the yellow fever mosquito, *Aedes aegypti* (Diptera: Culicidae)"

**OR6**

```
                                         (1)
CQUJHB000029_OR6       MYVFEDTLSSLNLMF-FASIPPCNEFYPETIRMIFKENKFFLIAFSLLLYTQLGEIVYLA	59
AAEL017548_OR6         MVSFEQSFTAVDLILITSGIPSCSSFYPVSIKSALKRNAVFLMAFTLLFYTAFGELVYLV	60
AALFPA_042417_OR6      MVKFEQTFKAVDLILITSGIPSCSSFYPASIKSALKRNAVFLIAFTLLLYTAFGELVYLV	60
                       *  **:::.:::*::  :.** *..*** :*:  :*.* .**:**:**:** :**:***.
                                                                           (0)
CQUJHB000029_OR6       QMVQRECSFLELTFQAPCVGYCSIGLLKMVVIAVKRNVIAELVQNLRDVWDQSIITVEHR	119
AAEL017548_OR6         EMLQRDFSFLEITFQAPCLGYCTIGLMKMLILAVKRNTIAELVQSLQEEWNKSVRSLEHQ	120
AALFPA_042417_OR6      QMLQREFTFLEITFQAPCLGYCSIGLMKMLILAMKRNSIAGLVQTLQKYWYTSVRSIEHQ	120
                       :*:**: :***:******:***:***:**:::*:*** ** ***.*:. *  *: ::**:

CQUJHB000029_OR6       EICKNVMKPAISITSLTAVVNVIMGISFTILPIAEMIYHYALAGGWVRQLPFLICWPFDP	179
AAEL017548_OR6         SICDDVMKPAIRFTTIVAVVNIVMGLAFTLLPIPEMIYYYSTHGKWVRQLPFLIWWSFDA	180
AALFPA_042417_OR6      SICDEVMKPAIRFTTIVAVVNIVMGMAFTLLPIPEMIYHYTITGQWVRQLPFLIWWSFDV	180
                       .**.:****** :*::.****::**::**:*** ****:*:  * ********* * **
                                               (0)
CQUJHB000029_OR6       LSGNKYFLVYPLYIVIGFSGIIIHMAFDCLFCILTAHICMNFRILEHDFENIKSASDINE	239
AAEL017548_OR6         YSGFVYYFIYPLYVVIGFSGIIIHMGFDCLFCILSAHLCVHLRILKFDLENLTYGLDSSE	240
AALFPA_042417_OR6      YAGFVYYFIYPLYIVIGFSGIIIHMGFDCLFCILAAHICVHFRILKHDLGNLTDLLDGTE	240
                        :*  *:::****:***********.********:**:*:::***:.*: *:.   * .*
 				      (2)                             (0)
CQUJHB000029_OR6       TS----E-SITKCLIHVEKHQR-LHYKEAVDGIFSFTLCYNFFVSSIIICIQGFMITAAS	293
AAEL017548_OR6         NM---SIKLNPKLFYIVEKHQNILECHDRMNKIFNFALFYNFFVSSFIICIQGFMVTAAS	297
AALFPA_042417_OR6      NIIENRKKVNPKLFGIVEKHQIIEECHNRMNTIFNFALFYNFFVSSFIICIQGFMVTAAS	300
                       .          * :  *****   . :: :: **.*:* *******:********:****
                                                     (0)
CQUJHB000029_OR6       GYTLIKFALFLASFLVELFLLCFYGQDVSESSSQVAQAAYNCSWYNESKKFRQLILQIIM	353
AAEL017548_OR6         GYTLIKFALFLASFLVELFLLCFYGHHIVESSVLVAEAAYNCLWYNTNHQFRTIILQMVN	357
AALFPA_042417_OR6      GYTLIKFALFLASFLVELFLLCFYGHHIVESSILAAEAAYHCSWYNTNHQFRTIVLQMIN	360
                       *************************:.: ***  .*:***:* *** .::** ::**::
				    (0)
CQUJHB000029_OR6       KAQRPLVLMAWKFWPVNIMTFSSILSASWSYFTLLRTVYYK	394
AAEL017548_OR6         KGQIPLSLMAWKIWPVNMNTFANILSASWSYFTLIRTVYAD	398
AALFPA_042417_OR6      KGQNPLTLMAWKIWPVDMSTFASILSASWSYFTLLRTVYAD	401
                       *.* ** *****:***:: **:.***********:**** .
```

**OR8**

```
AAEL012254_OR8         MNDLVKFESFIRVPEIFFDMIGITRYGEARDTWKARLKQAFFWSSYANTIFCLIIEHIYF	60
AALFPA_067679_OR8      MNDLVKFESFIRVPEIFFDMIGITRYGEPRDTWKARLKQAFFWSSYANTIFCLIIEHIYF	60
CQUJHB020308_OR8       MNDLVRFESFIRVPEIFYGMIGITRYGEPRNTTKARLKQLFFWSSYANTIFCLIIEHIYF	60
                       *****:***********:.********* *:* ****** ********************

AAEL012254_OR8         IKAAGNFTNFLELTALAPCIGFTALSIVKIMTIKLNEAKLNGILDRLSDLFPRSHLDQDR	120
AALFPA_067679_OR8      IKAAGNFTNFLELTALAPCMGFTALSIVKIMTIKLNEAKLNGILDRLSDLFPRSHQDQDR	120
CQUJHB020308_OR8       FRAAGNFTNFLELTALAPCIGFTALSIVKIMTIKLNEAKLNGILDRLKELFPVTHLEQTR	120
                       ::*****************:***************************.:*** :* :* *

AAEL012254_OR8         YRTYNYNLESQMVMKSFSILYMILIWIFNLLPLVSMLVNYISTGILEKELPYFMWYWYDW	180
AALFPA_067679_OR8      YRTYNYNLESQMVMKSFSILYMILIWIFNLLPLVSMLVNYLSTGVLEKELPYFMWYWYDW	180
CQUJHB020308_OR8       YRTHQYNLESQMVMKSFSILYMILIWIFNLLPLVSMLVNYLMSGVLVRELPYFMWYWYDW	180
                       ***::***********************************: :*:* :************

AAEL012254_OR8         HKAGYYEITFFHQNWGAFDSAVFNLSTDLLFCAIILLICLQFDILAYRLRHAKGDYKELE	240
AALFPA_067679_OR8      HKVGYYEITFFHQNWGAFDSAVFNLSTDLLFCAIILLICLQFDILAYRLRHAKGDYKELE	240
CQUJHB020308_OR8       HREGLYEITFFHQNWGAFDSAVFNLCTDLMFCAVILLMCLQFDIIAVRLRAAKDDPQELI	240
                       *: * ********************.***:***:***:******:* *** **.* :**

AAEL012254_OR8         QCVKLHQSVVELSNQLEGIFSPSILVNFVGSSVIICLVGFQATSNISAFDLFKFILFLIS	300
AALFPA_067679_OR8      ECAKLHQSVIELSNQLEGIFSPSILVNFVGSSVIICLVGFQATSDISAFDLFKFILFLIS	300
CQUJHB020308_OR8       SCVQLHQTVLELGDQLESIFSPSILVNFLGSSVIICLVGFQATSQISAFDLFKFVLFLIS	300
                       .*.:***:*:**.:***.**********:***************:*********:*****
                                        (0)
AAEL012254_OR8         SLVQVFLLCYYGNKLIEASSQIGYCAFEGTWYMADLRYQKSLLFVMTRAGQWQKLTAMKF	360
AALFPA_067679_OR8      SLVQVFLLCYYGNKLIEASSQITYCAFEGTWYMADVRYQKSLLFVMTRAGKWQKLTAMKF	360
CQUJHB020308_OR8       SLVQVFLLCYYGNKLIEASSQIPYAAFQGEWYLADVRYRKSLLFLMARAGKWQKLTAMKF	360
                       ********************** *.**:* **:**:**:*****:*:***:*********
                               (0)
AAEL012254_OR8         SVVSLASYSAILSTSFSYFTLLKTIYEPSQK-	391
AALFPA_067679_OR8      SVVSLASYSAILSTSFSYFTLLKTIYEPK---	389
CQUJHB020308_OR8       SVVSLASFTGILSTAFSYFTLLKTIYDPSDSK	392
                       *******::.****:***********:*.
```

**OR10**

```
AAEL006003_OR10         --MASILDCPIVSVNARVWRFWSFVLKHDAMRYISIIPVTVMTFFMFTDLCRSWGNIQEL	58
AALFPA_080165_OR10      --MESILNCPIVSVNARVWRFWSFVLKHDAMRYISIIPVTVMTFFMFLDLGHSWGDFQDV	58
CQUJHB000274_OR10       MTAAPILDCPIISVNVRVWHFWSFVLKHDAMRYISIIPVGVMNVFMFADLYRAWGNIDEV	60
                             **:***:***.***:******************* **..*** ** ::**:::::
```

(0) (0)

```
AAEL006003_OR10         IIKAYFAVLYFNAVLRTLILVKDRKLYENFMQGISNVYFEISHIDDHKIQSLLKSYTVRA	118
AALFPA_080165_OR10      IIKGYFAVLYFNAVLRTLILVKDRKLYENFMEGISKFYFEISRIDDHQIQSLLRSYTARA	118
CQUJHB000274_OR10       IINAYFAMIFFNAVLRTIFILCNRQDYEDFLQRIAEVYSEIAMIDDHVVQKLVRKFTKRA	120
                        **:.***:::*******:::: :*: **:*:: *::.* **: **** :*.*::.:* **

AAEL006003_OR10         RMLSISNLALGAIISTCFVVYPIFTGERGLPYGMFIPGLDSFRSPHYEIIYIVQVVLTFP	178
AALFPA_080165_OR10      RMLSISNLALGAIISTCFTVYPMFTGVRGLPYGMYIPGVDGYQSPQYEIIYLVQVVLTFP	178
CQUJHB000274_OR10       RLLSKANLVLGAVISTCYVVYPLFTGTRSLPYGMFIPGVNNFKTPLYQVFFIGQAVLTFP	180
                        *:** :**.***:****:.***:*** *.*****:***::.:::* *::::: *.*****

AAEL006003_OR10         GCCMYIPFTSFFASTTLFGLVQIKTLQRQLQTFKDNISSQDKEKVKTKVVKLIEDHKRII	238
AALFPA_080165_OR10      GCCMYIPFTSFFVSTTLFGLVQIKTLQRQLQTFKDGIGSHGNKNANLQVIKLIQDHKRII	238
CQUJHB000274_OR10       GCCMYIPFTSFFATTTLFGLVQIQTLQRQLRTFKDEVVKENRALVESKLEKCIEDHKRII	240
                        ************.:*********:******:**** : ....  .: :: * *:******
                                                         (0)
AAEL006003_OR10         TYVSELNSLVTYICFVEFLSFGMMLCALLFLLNVIENHAQIVIVAAYIFMIISQIFAFYW	298
AALFPA_080165_OR10      AYVSELNSLVTYICFVEFLSFGLMLCALLFLLNVIENHAQMVIVAAYIFMIISQIFAFYW	298
CQUJHB000274_OR10       RYVSDVNSLVTYICLIEFMSFGLMLCALLFLLNIIENPAQIIIVVAYIFMIISQIFTFYW	300
                         ***::********::**:***:**********:*** **::**.***********:***
                                                                   (0)
AAEL006003_OR10         HANEVREESMNLAEAAYSGPWVELDNSIKKKLLLIILRAQQPLEITVGNVYPMTLEMFQS	358
AALFPA_080165_OR10      HANEVREESMNIAEAAYSGPWVELDDSIKKKLLLIILRAQQPLEITVGNVYPMTLEMFQS	358
CQUJHB000274_OR10       HANELREESMGIAEAAYDAPWVELDDSMKKKLLLIIARAQQPLEITVGNVYAMTLEMFQS	360
                        ****:*****.:*****..******:*:******** ************** ********

AAEL006003_OR10         LLNASYSYFTLLRRVYN	375
AALFPA_080165_OR10      LLNASYSYFTLLRRVYN	375
CQUJHB000274_OR10       LLNASYSYFTLLRRVYN	377
                        *****************
```

**OR11**

```
AAEL011583_OR11          MQLKDEWIQEDDVYDNPLLRLTINGLKYYGILLYKSQPFKKLNCFRGVCFTASMLAFNVT	60
AALFPA_064092_OR11a      MQLKDEWIQEDDVYDNPLLRLTINGLKYYGILLYKSQPFKKLNCFRGVCFTASMLAFNVT	60
AALFPA_060890_OR11b      MQLKDEWIQEDDVYDNPLLRLTINGLKYYGILLYKSQPFKKLNCFRGVCFTASMLAFNVT	60
CQUJHB004338_OR11        MELKDEWIQEDDVYSNPLLRLTLNGLKYYGILLYKSQPLKKLNCFRGVCFTASMLAFNVT	60
                         *:************.*******:***************:*********************
                                                                 (1)
AAEL011583_OR11          QYVDLYQVWGNIAEMTANAATTLLFTTTIVRILHFYWNRARFNNAIKVADEGVQHLLRFG	120
AALFPA_064092_OR11a      QYVNLYQVWGDISQMTANAATTLLFTTTIVRILHFYWNRARFNNAIKVADEGVQHLLRFG	120
AALFPA_060890_OR11b      QYVNLYQVWGDISQMTANAATTLLFTTTIVRILHFYWNRARFNNAIKVADEGVQHLLRFG	120
CQUJHB004338_OR11        QYVDLVQVWGDIGEMTANAATTLLFTTIILRIFHFYWNRARFNNAIKFADEGVRHILDYG	120
                         ***:* ****:*.:************* *:**:**************.*****:*:* :*

AAEL011583_OR11          NAPEKEIFWDNVKYMNRLTAAFWICALVTANTMCVYALVQYQSLKSMDSFNSTEPFDPPT	180
AALFPA_064092_OR11a      NAPEKAIFWDNVRYMNRLTAAFWICALVTANTMCVYSLVQYHSLKSMEPFNSTKPFDPPT	180
AALFPA_060890_OR11b      NAPEKAIFWDNVRYMNRLTAAFWICALVTANTMCVYSLVQYHSLKSMEPFNSTKPFDPPT	180
CQUJHB004338_OR11        TPAEKAIFWDNVGYMKRLTAVFWICALVTANTMCVYALIEYCSVP------EAERVEPPM	174
                         .  ** ****** **:****.***************:*::* *:       .:: .:**

AAEL011583_OR11          ILRSWYPTDNIVDSFATIYLIQLYIMYVGQLIVPCWHVFMVSLMLYARTALMALNYKLAN	240
AALFPA_064092_OR11a      ILRSWYPTDNIMESFTTIYLIQLYIMYVGQLIVPCWHVFMVSLMLYARTSLMVLNYKLAN	240
AALFPA_060890_OR11b      ILRSWYPTDNIMESFTTIYLIQLYIMYVGQLIVPCWHVFMVSLMLYARTSLMILNYKLAN	240
CQUJHB004338_OR11        ILRSWYPGGHKEENFGAIYAVQLYIMYVGQLIVPCWHVFIVSLMVYVRAALTILNHKLRH	234
                         ******* .:  :.* :** :******************:****:*.*::*  **:** :

AAEL011583_OR11          LEQYAVSGMRSGRKIKCVVDPEEERCNVRKELIVECIQQQHKIFEYTRELEALTRGAMFM	300
AALFPA_064092_OR11a      LEMYAVSGLRSGRKLKLVKDPEEERCNERKELIVECIQQQHKIFEYTRELEALTRGAMFM	300
AALFPA_060890_OR11b      LEMYAVSGMRSGRKLKLVKDPEEERCNERKELIVECIQQQHKIFEYTRELEALTRGAMFM	300
CQUJHB004338_OR11        LDAYVTSGMRSGRLIRELKDPEEERLNARRELIIECVQRQRKLHEYTDELESLIQGPVFL	294
                         *: *..**:**** :: : ****** * *:***:**:*:*:*:.*** ***:* :* :*:
                                          (0)                               (0)
AAEL011583_OR11          DFVVFSVLLCALLFEASSTNSFVQIFIDICYIMTMTAMLFLYYWHANEIHYQANLLSSSA	360
AALFPA_064092_OR11a      DFVVFSVLLCALLFEASSTDSFVQIFIDICYIMTMTAILFLYYWHANEIHYHANLLSSSA	360
AALFPA_060890_OR11b      DFVVFSVLLCALLFEASSTDSFVQIFIDICYIMTMTAILFLYYWHANEIHYHANLLSSSA	360
CQUJHB004338_OR11        DFIVFSVLLCALLFEASVTDSAVQVFIDVCYIMTMTAILFLYYWHANEIQHQSDQLSKSA	354
                         **:************** *:* **:***:********:***********::::: **.**
                                                                  (0)
AAEL011583_OR11          FMNDWYNYPRSVNRHLITFICYSNKPLDMKAYIVSMSLDTFLAILRASYSYFTILKQAAG	420
AALFPA_064092_OR11a      FMNDWYNYPRSVNRHLITFICYSNKPLNMKAFIVSMSLDTFLAILRASYSYFTILKQAAG	420
AALFPA_060890_OR11b      FMNDWYNYPRSVNRHLITSICYSNKPLNMKAFIVSMSLDTFLAILRASYSYFTILKQAAG	420
CQUJHB004338_OR11        FANDWYNYPPKVNRNLLVLICYSIKPRIMKAFIVSMSLDTFIAILRASYSYFTILKQAAD	414
                         * ******* .***:*:. **** **  ***:*********:*****************.
```

**OR13**

```
AAEL008368_OR13           MWQPLRKFLAPGPELLSFGLQMLRFIGLWGDRRQVVRYLLVLFSEFIFLIGPKALLGSDK	60
AALFPA_046408__OR13a      MWPRLRKFLAPEPELLSFGLQMLRFVGLWGDRRRIVRYLLVLLSELMFLIGPKALLGSGK	60
AALFPA_064397_OR13b       MWPRLRKFLAPEPELLSFGLQMLRFVGLWGDRRRIVRYLLVLLSELMFLIGPKALLGSGK	60
CQUJHB000018_OR37         ----MSPAIPPDLQVLKFPLRMLRFVGLWGDRRELVRYASVVLCMSVVLIIPKAALGSGK	56
                              :   : *  ::*.* *:****:*******.:***  *::.  :.** *** ***.*

AAEL008368_OR13           EGFDSTARNIGELIFLVEVCISIGIFASRRASFERLIVVLENILRRKWPRNLQDEIYRFH	120
AALFPA_046408__OR13a      EGIDSTVRNIAELIFLVEVCISIGIFASRRTSFERLMAILEEILRRKWPQDLQDEIDRFH	120
AALFPA_064397_OR13b       EGIDSTVRNIAELIFLVEVCISIGIFASRRTSFERLMAVLEEILRRKWPQDLQDEIDRFH	120
CQUJHB000018_OR37         DGFDSFARNTAELIFFTEVCVSIGIFASRRGSFERLVEVLRETVLMYEDVELLGEIAAFN	116
                          :*:** .** .****:.***:********* *****: :*.: :      :* .**  *:
                                                                           (2)
AAEL008368_OR13           RRMEFFARAYALYIGFLLFLYNCVPIGSTIVKLIRFDESERSDFMLVVELQFFWFDIRRN	180
AALFPA_046408__OR13a      RRMEFFARAYALYIGFLVVLFCCVPVGSTLVKLVRFDESERSDFMLVVELQFFWFDIRRN	180
AALFPA_064397_OR13b       RRMEFFARAYALYIGFLVVLFCCVPVGSTLVKLVRFDESERSDFMLVVEL-FFWFDIRRN	179
CQUJHB000018_OR37         RKMERFSKSYAAWIGFWVVLYLGIPMIFTCVKVVFPGEGDRGDFMLIAELQFYWLDIRRN	176
                          *:** *:::** :*** :.*:  :*:  * **::  .*.:*.****:.** *:*:*****

AAEL008368_OR13           VVHYAIYMAFCFVAVSCSAYQSTLKGSVIVVVTQYGSKLFELISKRIDAMRSIERAADRD	240
AALFPA_046408__OR13a      AFHYAIYMMFCLVAVSCSAYQSVLKGSIIVVVTQYGSKLFELISKRIEAMAKIPKQTDRD	240
AALFPA_064397_OR13b       AFHYAIYMMFCLVAVSCSAYQSVLKGSIIVVVTQYGSKLFELISKRIEAMAKIPKQTDRD	239
CQUJHB000018_OR37         LLDYAIYLVFCSMAIFCSSYQSTLKGAVILVSIQYGTKLFELVAMSIDRLGNVKEEIARK	236
                           ..****: ** :*: **:***.***::*:*  ***:*****::  *: : .: .   *.
	 (0)
AAEL008368_OR13           RELREIVKLHGMALEYVQHLESTISFVMINQIMNCIFIWCLMMFYVSTNFGPNAANVMLL	300
AALFPA_046408__OR13a      RELREIVKLHGLAMEYVHHLESTISFVMINQIMNCILIWCLMMFYVSTNFGPNAANVMLL	300
AALFPA_064397_OR13b       RELREIVKLHSLAMEYVHHLESTISFVMINQIMNCILIWCLMMFYVSTNFGPNAANVMLL	299
CQUJHB000018_OR37         NQLREIVNLHKLAFQYTKHLEDTVCFMMINQILNCILIWCLMMFYVSTNFGPNAVCVIIL	296
                          .:*****:** :*::*.:***.*:.*:*****:***:*****************. *::*
                                              (0)
AAEL008368_OR13           FLVLMGEMVVYCLNGTALSEQAAGVGHAIYNYPWYKESVAMQKNMQLMIQRAQRPTGITA	360
AALFPA_046408__OR13a      FLVLMGEMVVYCLNGTALSEQAAGVSHAIYHYPWYTESVQMQKHMQLIIQRAQRPTGVTA	360
AALFPA_064397_OR13b       FLVLMGEMVVYCLNGTALSEQAAGVSHAIYHYPWYTESVQMQKHMQLIIQRAQRPTGVTA	359
CQUJHB000018_OR37         FAVLMGEMIAYCVNGSKLAETAAAVGHAVYRYPWYNEPTAMQKDMQLIIERAQKPTGITA	356
                          * ******:.**:**: *:* **.*.**:*.****.* . ***.***:*:***:***:**

AAEL008368_OR13           AKFYFVNIERLGLVTQASYSYYLILKNRF	389
AALFPA_046408__OR13a      AKFYYVNIERLGMVIQASYSYYLILKKRF	389
AALFPA_064397_OR13b       AKFYYVNIERLGMVIQASYSYYLILKKRF	388
CQUJHB000018_OR37         AKFYFVNIERLGLVVQASYSYYLILKKRF	385
                          ****:*******:* ***********:**
```

**OR15**

```
AAEL008448_OR15         MKYFELTEPEAAMPLALRLLETYGLRGGKRKFLQFQVTILWELLMIVIPKIVFGYRS-QD	59
AALFPA_076765_OR15      MKYFELVEPEAVMPLALRLMETYGLRGGKRKFLQFQVTILWEFLTIVLPKIIFGYRS-QD	59
CQUJHB006521_OR1        MKFAPLQNRMAVMPFTLQYLRLFGLRGDRRNRVHFVLALLYRVLLLNFPKLAFGFRDRID	60
                        **:  * :  *.**::*: :. :****.:*: ::* :::*:..* : :**: **:*.  *
                                                                  (1)
AAEL008448_OR15         LVIRGLSELLFQLHIMIRISIFAWHRFKYESLIDIIRKVYRKTFSTGGDPTSKSIILKFN	119
AALFPA_076765_OR15      LVIRGLSELLFQLHIMIRISIFAWHRFKFEDLVAIIRRVYRKTFSPGADSKLKEIILDFN	119
CQUJHB006521_OR1        LVIRSISELLFQIHIDLRAVLFATKLREFEELAGLLRKVYNKVKTLDADSPERKIIEASN	120
                        ****.:******:** :*  :** :  ::*.*  ::*:**.*. : ..*   :.**   *
	(2)
AAEL008448_OR15         QMINKQSKGYFLYIMGCVSLFSVAPVVQSVIIFMANQSRNGTEKAEYVTMMEQEFYGLDI	179
AALFPA_076765_OR15      KMINKQSKGYFLYIMGCVSLFTVAPVVQSVVIFITHQGNNGTDKAEYVTMMEQEFYGLNI	179
CQUJHB006521_OR1        LGINRRSKSYALYVAIAVTVFFWVPVVQTTAIWLLNRGSNSTDRPEFVTMMELEFYGLDI	180
                          **::**.* **:  .*::*  .****:. *:: ::. *.*:: *:***** *****:*

AAEL008448_OR15         RGNFGHYAIYVALAGLAHYYSASFFAVTGVIIICGVRCTILTFKLINVRLSKLHELPKQD	239
AALFPA_076765_OR15      RGNFAHYVMYVGLAGLAHYYSASFFAVTGVVMICAVRCTILTFRLIIVRLSKLHELPKED	239
CQUJHB006521_OR1        RGNIWHYLVYASLSSVAHYYSAVYFALSGMVIFSCIKSIAALFELVSTRLATLHELSGKE	240
                        ***: ** :*..*:.:****** :**::*::::. ::.    *.*: .**:.****  ::
                                        (2)                              (0)
AAEL008448_OR15         IRDELREIIDLHVDALRCIQLLEQIANLAMVIQIIDCVLIWISMILYMRNNLGVDAISLM	299
AALFPA_076765_OR15      IRGELREIIDLHVDALRCIQLMEQIANLAMVIQIVDCVLIWISMILYMRNNLGVDAISLM	299
CQUJHB006521_OR1        LREELADLVELHVNGLRCIELLENINNLAMMVQMVNCVLIWISMFLSISTNFTPEVVSLL	300
                        :* ** ::::***:.****:*:*:* ****::*:::********:* : .*:  :.:**:
                                             (0)
AAEL008448_OR15         VLFVALTGETYALCDLLTQLTSESLAVTRAIIDCQWYSLPLDVQKSLSFVLFRAQRKEGI	359
AALFPA_076765_OR15      VLFVALTGETYALCDLLTQLTSESLAVTRAIIDCQWYNLPLDVQKALSFILFRAQRNEGI	359
CQUJHB006521_OR1        VLLIVMTGETYVLCQLATELSHVNLTVAESIHRSEWIQMPVDVQKGLAMMLQRAQKREGL	359
                        **::.:*****.**:* *:*: ..*:*:.:*  .:* .:*:****.*:::* ***:.**:

AAEL008448_OR15         TAAKFFFMDIERFGSVAQTSYSIYVVLKDQL	390
AALFPA_076765_OR15      TAAKFFYMDIERFGNVAQTSYSIYVVLKDQL	390
CQUJHB006521_OR1        TAAKFFYMDVERFGRVAQTSYSIFIVLKERI	390
                        ******:**:**** ********::***:::
```

**OR28**

No homologues

**OR31**

```
AAEL013217_OR31          --------------------------MAPTQNGRDREKFLRVQLLCLALIGIKRHETVSS	34
AALFPA_070001_OR31a      -------------------MALPSPTPADGRRTTSQGKFLRVQLLCLALIGVQRLETVSS	41
AALFPA_067456_OR31b      -------------------MALPSPTPADGRRTTSQGKFLRVQLLCLALIGVQRLETVSS	41
AALFPA_062046_OR31c      -------------------MALPSSTPVDGRTTTAQGKFLRVQLLCLALIGVQRDETVSS	41
CQUJHB008764_OR95        MA---------TFTSKIAWTANDGNHRRGRSPDKEKDNFLRVQLFTMALNGIRKHETVSS	51
CQUJHB011798_OR96        MATTTAKATSNPGTPRAAWVS-----DVGGRTIDDEENFLRVQLFSMALIGVQERETVPS	55
                                                            . :******: :** *::. *** *

AAEL013217_OR31          RTIFHVCFISMVIMDLATILFALEHANDIALVCDCLGPTFTAYLGIVKQYCLSAHRVELW	94
AALFPA_070001_OR31a      RAIFHVCFISMVIMDLATILFALEHANDIALVCDCLGPTFTAYLGIVKQYCLSAHRVELW	101
AALFPA_067456_OR31b      RAIFHVCFISMVIMDLATILFALEHANDIALVCDCLGPTFTAYLGIVKQYCLSAHRVELW	101
AALFPA_062046_OR31c      RAIFHVCFISMVIMDLATILFALEHANDIALVCDCLGPTFTAYLGIVKQYCLSAHRVELW	101
CQUJHB008764_OR95        RIYFYGCFLTMLIMDLAGVHFAYQNAGEILLVCDCLGPTFTCFLGVVKQYYLDVHREQLW	111
CQUJHB011798_OR96        RIYSYFCFYSLLIMDLSMVLFAVQHFGDMVLVCDCLGPGFTAYLGMVKQHYLSEQRKQLW	115
                         *   : ** :::****: : ** :: .:: ******** **.:**:***: *. :* :**
	      (1)
AAEL013217_OR31          NIIETLRRLKDYAGTSEIESIERNNKIDRFLATAYLMSASATGSLFIIAALAKGCYKLIF	154
AALFPA_070001_OR31a      NIIETLRRLKDYAQADEIVSIERNNKIDRILATAYLVSASATGSLFIFAALAKGFYKLIF	161
AALFPA_067456_OR31b      NIIETLRRLKDYA--DEIVSIERNNKIDRILATAYLVSASATGSLFIFAALAKGFYKLIF	159
AALFPA_062046_OR31c      NIIETLRRLKDYAQADEIVSIERNNKIDRILATAYLVSASATGSLFIFAALAKGFYKLIF	161
CQUJHB008764_OR95        YIIHELRKLKQNATVSDIEMIEKNNKIDQFLATAYFASSMGTGTIFIVEAILKGTYNYFI	171
CQUJHB011798_OR96        EIIYALKKLKQIAKPDEIRSIERNNQIDRYLATAYLTSAVITGSHFIVTAIVKAVYSKVV	175
                          **  *::**: *  .:*  **:**:**: *****: *:  **: **. *: *. *. ..
	 (2)
AAEL013217_OR31          QNIIEWGFPLSLSFPFKTSHPIVFGVFFVWSSAAIYIVVFCSVSSDASFGGLASNVVVHF	214
AALFPA_070001_OR31a      QNSIEWGLPLSLSFPFNTSQPIIFALFFIWSSAAIYIVVFCSVSSDASFGGLASNVVVHF	221
AALFPA_067456_OR31b      QNSIEWGLPLSLSFPFNTSQPIIFALFFIWSSAAIYIVVFCSVSSDASFGGLASNVVVHF	219
AALFPA_062046_OR31c      QNSIEWGLPLSLSFPFNTSQPIIFALFFIWSSAAIYIVVFCSVSSDASFGGLASNVVVHF	221
CQUJHB008764_OR95        RNCIEWNLPIAISFPFDISHPAIFAFFFIWCSAATYMVVFSSVSSDAGFGGLASNLVVHF	231
CQUJHB011798_OR96        HGKFVWQLPLLQSYPFDISHPLMFAVLFVWTSATIYMVVFGSVSSDAAFGGLASNLVVHF	235
                         :. : * :*:  *:**. *:* :*..:*:* **: *:*** ******.*******:****

AAEL013217_OR31          KLLQKRLQDATFADNDENLKQLIEYHSLLLNLSRKIMSSFRVIIINNLLVASVLLCVLGF	274
AALFPA_070001_OR31a      KLLQKRLQDTTFTDNDENLKYLIEYHSLLLGLSRKLMSSFRLIIINNLLVASVLLCVLGF	281
AALFPA_067456_OR31b      KLLQKRLQDTTFTDNDENLKYLIEYHSLLLGLSRKLMSSFRLIIINNLLVASVLLCVLGF	279
AALFPA_062046_OR31c      KLLQKRLQDTTFTDNDENLKYLIEYHSLLLDLSQKLMSSFRVIIINNLLVASVLLCVLGF	281
CQUJHB008764_OR95        KILQNRFKDRRFEDDDQSLEDLIEYHTLVLKLSRRLMSSFRIIILNNLLVASVLLCVLGF	291
CQUJHB011798_OR96        KFIQAGFRDRSFEDNDPSLKDLIEYHRHVLDLSRKLISAYRPIMLNNFIVASFLLCVLGF	295
                         *::*  ::*  * *:* .*: *****  :* **::::*::* *::**::***.*******
	(0)
AAEL013217_OR31          QLVMFLGSTLMLIYLMYVTAIVIQITFFAYYGSLLSHESEEVSSSIYCSNWYEASPKTRR	334
AALFPA_070001_OR31a      QLVMFLGSTLMLIYFMYVTAIVIQITFFAYYGSLLSHESEAVGNAIYCSNWYEATPKTRR	341
AALFPA_067456_OR31b      QLVMFLGSTLMLIYFMYVTAIVIQITFFAYYGSLLSHESEAVGNAIYCSNWYEATPKTRR	339
AALFPA_062046_OR31c      QLVMFLGSTLMLIYFMYVTAIVIQITFFAYYGSLLSHESEAVGNAIYCSNWYEATPKTRR	341
CQUJHB008764_OR95        EMVVYLGTSLMLLYITYITAIVIQIFFFSYYGSQLLYESAAVGDAIYCSNWYEATPKTRK	351
CQUJHB011798_OR96        QLVLFMGSTMMFLYIVFVTAIVIQITFFSYYGSQLSHESALVGDAIYCSNWYETSPKTRR	355
                         ::*:::*:::*::*: ::******* **:**** * :**  *..:********::****:
    (0)
AAEL013217_OR31          ILLQCLMRAQVPVNTKAGFMVASLPTLRAILNSAGSYVALLLSFTDN	381
AALFPA_070001_OR31a      ILLQCLMRAQVPVNTKAGFMVASLPTLRAILNSAGSYVALLLSFTD-	387
AALFPA_067456_OR31b      ILLQCLMRAQVPVNTKAGFMVASLPTLRAILNSAGSYVALLLSFTD-	385
AALFPA_062046_OR31c      ILLQCLMRAQVPVNTKAGFMVASLPTLRAILNSAGSYVALLLSFTD-	387
CQUJHB008764_OR95        LLLFCIKRAQVPVNTKVGIMVASLPTFRAIVNSAGSYVALLLSFTED	398
CQUJHB011798_OR96        LLLQCLMRAQVPVNIRVGFIEASMPTFRAILNSAGSYVALLLSFTDT	402
                         :** *: ******* :.*:: **:**:***:**************:
```

**OR55**

No homologues

**OR71**

```
AAEL017564_OR71         MELSYHRSLPPELQIMPFQLRCMELIGLIGPKGRFYRFVLAFGWGTFVILLPKSVLGIGS	60
AALFPA_064397_OR71      MELSHYRSLPSELQIMPFQLRCMELIGLAGPKGRFYRFVLAFGWGTFVILLPKSVLGIGS	60
                        ****::**** ***************** *******************************
                                                                     (1)
AAEL017564_OR71         SELDAIIKGFAELLFEGNLFIAVASLVPKLPLVKRLLHVLSEIFRQATHDKRDVKDHCYA	120
AALFPA_064397_OR71      PRLDAIIKGFAELLFEGNLFIAVASLIPKLPLLKRLLHVLSEIFTQVTRDAR-SKDQCYE	119
                         .************************:*****:*********** *.*:* *  **:**

AAEL017564_OR71         LICEQNSKIDKFCKFYFIYCCFGPFVFCIPAMVTSYVRYFGTTNNGTNA---NGSEHLRF	177
AALFPA_064397_OR71      LICEQNSKIDKFCKFYFIYCCFGPFVFCIPAMVTSYVRYFGGAGNGTDNADSNETEQLLF	179
                        ***************************************** :.***:    * :*:* *
	(2)
AAEL017564_OR71         ELPMEQEFYWLPIRTNFACYHLFLTLSLSAYCVCSYMSVIKVSTLLIMIKYCSLVYRLVA	237
AALFPA_064397_OR71      ELPMEQEFYGLQIRTNFAHYHLFLAASLSAYCVCSYMSVIKVSTLLIMIKYSSLAYRLVA	239
                        ********* * ****** *****: *************************.**.*****
```

(0) (0)

```
AAEL017564_OR71         IRIRELGKLPPGRKQDD----TDEDERTKMVKVKEVVEMHEKALEATDLVEKVINIPIAM	293
AALFPA_064397_OR71      IRIRKLAELPASGQEQNFEEVTDGGDVLKMSMVKEVVDLHKKALEVTDLVEEIINIPIAM	299
                        ****:*.:** . ::::    ** .:  **  *****::*:****.*****::*******
                                         (0)
AAEL017564_OR71         QFMACILFWCMTMVYVSTNINFNLFNVMVLFWLSLIETYGYSYLGTELSEDAKAVGHAVY	353
AALFPA_064397_OR71      QFIACILFWCMTMFYVSTNINFNLFNVMVLFWLSLIETYGYSYLGTELTEEAKTVGQVIY	359
                        **:**********.**********************************:*:**:**:.:*

AAEL017564_OR71         DLPWYEDSAQLQRYYRLMIQRSQQNIGVTAAKFFIVGIEKFGKVVNLSYSYYLVLKDVLD	413
AALFPA_064397_OR71      ELPWYEDSAQLQRYYRLMIQRTQQNTGVTAAKFFIVGIEKFGKVVNLSYSYYLVLKDVLD	419
                        :********************:*** **********************************

AAEL017564_OR71         RL	415
AALFPA_064397_OR71      SL	421
                         *
```
